# S100A4 exerts robust mucosal adjuvant activity for co-administered antigens in mice

**DOI:** 10.1101/2022.01.04.474916

**Authors:** Arka Sen Chaudhuri, Yu-Wen Yeh, Jia-Bin Sun, Olifan Zewdie, Tao Jin, Bin Wei, Jan Holmgren, Zou Xiang

## Abstract

The lack of clinically applicable mucosal adjuvants is a major hurdle in designing effective mucosal vaccines. We hereby report that the calcium-binding protein S100A4, which regulates a wide range of biological functions, is a potent mucosal adjuvant in mice for co-administered antigens, including the SARS-CoV-2 spike protein, with comparable or even superior efficacy as cholera toxin but without causing any adverse reactions. Intranasal immunization with recombinant S100A4 elicited antigen-specific antibody and pulmonary cytotoxic T cell responses, and these responses were remarkably sustained for longer than six months. As a self-protein, S100A4 did not stimulate antibody responses against itself, a quality desired of adjuvants. S100A4 prolonged nasal residence of intranasally delivered antigens and promoted migration of antigen-presenting cells. S100A4-pulsed dendritic cells potently activated cognate T cells. Furthermore, S100A4 induced strong germinal center responses revealed by both microscopy and mass spectrometry, a novel technique for measuring germinal center activity. In conclusion, S100A4 may be a promising adjuvant in formulating mucosal vaccines, including vaccines against pathogens that infect via the respiratory tract, such as SARS-CoV-2.

## Introduction

The COVID-19 pandemic, caused by the virus SARS-CoV-2, continues to be a major global health menace. As vaccination is considered the most cost-effective strategy for the control of infectious diseases, strategies to control the COVID-19 pandemic by developing effective vaccination modalities have been intensely developed worldwide. To date the vast majority of available vaccines for human use are needle injection-based, with a few exceptions for a few licensed mucosal vaccines that include oral vaccines for polio, cholera, typhoid, and rotavirus diarrhea, as well as an intranasal vaccine for influenza *(Miquel-Clopes et al., 2019)*. Likewise, all the SARS-CoV-2 vaccines currently liscenced for human use are injection-based *(Bok et al., 2021)* despite the fact that SARS-CoV-2 accesses the body through the mucosal membranes of mainly the respiratory tract, at which sites the mucosal adaptive immune responses for preventing pathogen entry are in general more efficiently induced by topical-mucosal rather than parenteral vaccination *(Su et al., 2016)*. In particular, nasal immunization is effective for inducing immune responses in the respiratory tract *(Lobaina Mato, 2019)*. Moreover, mucosal vaccination obviates the use of needles, which is suitable for mass vaccination during pandemics such as COVID-19.

However, a hurdle for a wider use of mucosal vaccination is that with most purified antigens, successful immunization will usually require co-administration with an effective mucosal adjuvant. There is, as yet, a scarcity of safe and effective mucosal adjuvants for human use. Cholera toxin (CT) is the most potent and best characterized “gold standard” mucosal adjuvant used in experiments on animals but is too toxic for humans. We have recently identified a critical role for the calcium-binding protein S100A4 in the mucosal adjuvanticity of CT by demonstrating that endogenous S100A4 is required for CT’s ability to adjuvant both humoral and cellular adaptive immune responses following mucosal immunization *(Sun et al., 2017)*. Mice that were genetically deficient in S100A4 mounted severely compromised antigen-specific immune responses, which were associated with dampened antigen presentation by dendritic cells (DCs), reduced T cell activation, and a lack of germinal center formation. Notably, the engraftment of wild-type DCs in the S100A4-deficient mice prior to mucosal immunization with the antigen, together with CT, almost fully restored the defective immune responses, indicating that the expression of S100A4 by DCs and possibly other antigen-presenting cells is critical for mounting an effective adaptive immune response *(Sun et al., 2017)*.

S100A4 is one of more than 20 calcium-binding proteins collectively categorized as the S100 protein family *(Gross et al., 2014)*. S100A4 was originally described as fibroblast-specific protein 1 (FSP1) because of its restricted expression to stromal fibroblasts *(Strutz et al., 1995)*. However, this protein was later found to have a much wider expression spectrum, being expressed also by at least endothelial cells, smooth muscle cells, lymphocytes, neutrophils, macrophages, and DCs *(Boomershine et al., 2009, Li et al., 2010, Mishra et al., 2012)*. S100A4 interacts with cellular targets through at least two receptors: the receptor for advanced glycation end-products (RAGE) and toll-like receptor 4 (TLR4) *(Boye et al., 2008, Cerezo et al., 2014)*. S100A4 has the capacity to regulate a diverse range of cellular processes, such as cell growth, survival, differentiation, and motility *(Donato et al., 2013)*. We have previously revealed a critical role of S100A4 in adaptive immune responses *(Bruhn et al., 2014, Sun et al., 2017)*.

In this study, we examined the potential of using purified exogenous S100A4 as a mucosal adjuvant in its own right for co-administered antigens. Our results demonstrate that when given intranasally, together with either ovalbumin (OVA), as an experimental vaccine antigen, or the spike protein of SARS-CoV-2, as a clinically applicable vaccine antigen, S100A4 can, without any adverse reactions, effectively augment antigen-specific humoral immune responses both mucosally and systemically, as well as stimulate antigen-specific cytotoxic T cell responses in the lungs, to levels comparable to or better than those using CT as adjuvant. Remarkably, these antigen-specific adaptive immune responses were sustained for longer than six months. We also introduced a novel, label-free matrix-assisted laser desorption/ionization (MALDI) time-of-flight (TOF) mass spectrometry (MS) technique for the detection of S100A4-induced germinal center responses in the spleen.

## Results

### Exogenous S100A4 is a strong mucosal adjuvant

Based on our previous findings of endogenous S100A4 being critical for the *in vitro* antigen-presenting capacity of DCs, and for the *in vivo* mucosal adjuvant effect of CT *(Sun et al., 2017)*, we wished to examine whether exogenous S100A4 might also work as a mucosal adjuvant in its own right. We tested this by immunizing mice intranasally on three occasions with a model antigen, OVA, administered with or without S100A4, and for comparison in a separate group of mice also with CT, and by examining immune responses three days after the last dose (Supplementary Fig. 1a). Serum anti-OVA total IgG, IgG1, and IgG2c levels were all robustly augmented after immunization together with S100A4, to levels comparable to those achieved using CT as adjuvant (Supplementary Fig. 1b). S100A4 also potentiated mucosal antibody production in mucosal tissues, resulting in strongly enhanced OVA-specific IgA, total IgG, and IgG1 antibody levels in lung exudate (Supplementary Fig. 1c). Importantly, the intranasal co-administration with S100A4 further resulted in enhanced antigen-specific IgA and IgG responses at a remote site, the vaginal mucosal surface (Supplementary Fig. 1d), consistent with previous studies showing a nasal-genital tract mucosal immunological link *(Czerkinsky & Holmgren, 2010)*. Similar to the serum antibody responses, the immunization with S100A4 also promoted mucosal antigen-specific antibody production to levels comparable to those achieved by using CT as adjuvant (Supplementary Fig. 1).

Next, we largely repeated this experiment but halved the amount of S100A4 (10 μg per dose) and extended the antibody measurement to as long as 196 days (roughly 6.5 months) after the last immunization (Fig. 1a). At this substantially late time point, serum anti-OVA IgG antibody responses, including total IgG, IgG1, and IgG2c, remained high in S100A4-adjuvanted mice, not much less compared to a much earlier time point, 10 days after the last dose of immunization (Fig. 1b). Furthermore, mucosal anti-OVA IgG and IgA levels, in lung exudate (Fig. 1c), bronchoalveolar lavage fluid (BALF) (Fig. 1d), vaginal lavage (Fig. 1e), and feces (Fig. 1f), remained substantially high at this late time point in mice which received OVA admixed to S100A4. The S100A4-driven antibody sustainability was comparable to CT-adjuvanted immunization (Fig. 1b–f). These late time point mucosal antibody levels were also found to be sustained in comparison to the levels in previous experiments measured at an earlier time point (Supplementary Fig. 1c and d), despite a higher dose of S100A4 being used in that immunization (20 μg per dose).

**Figure 1.**
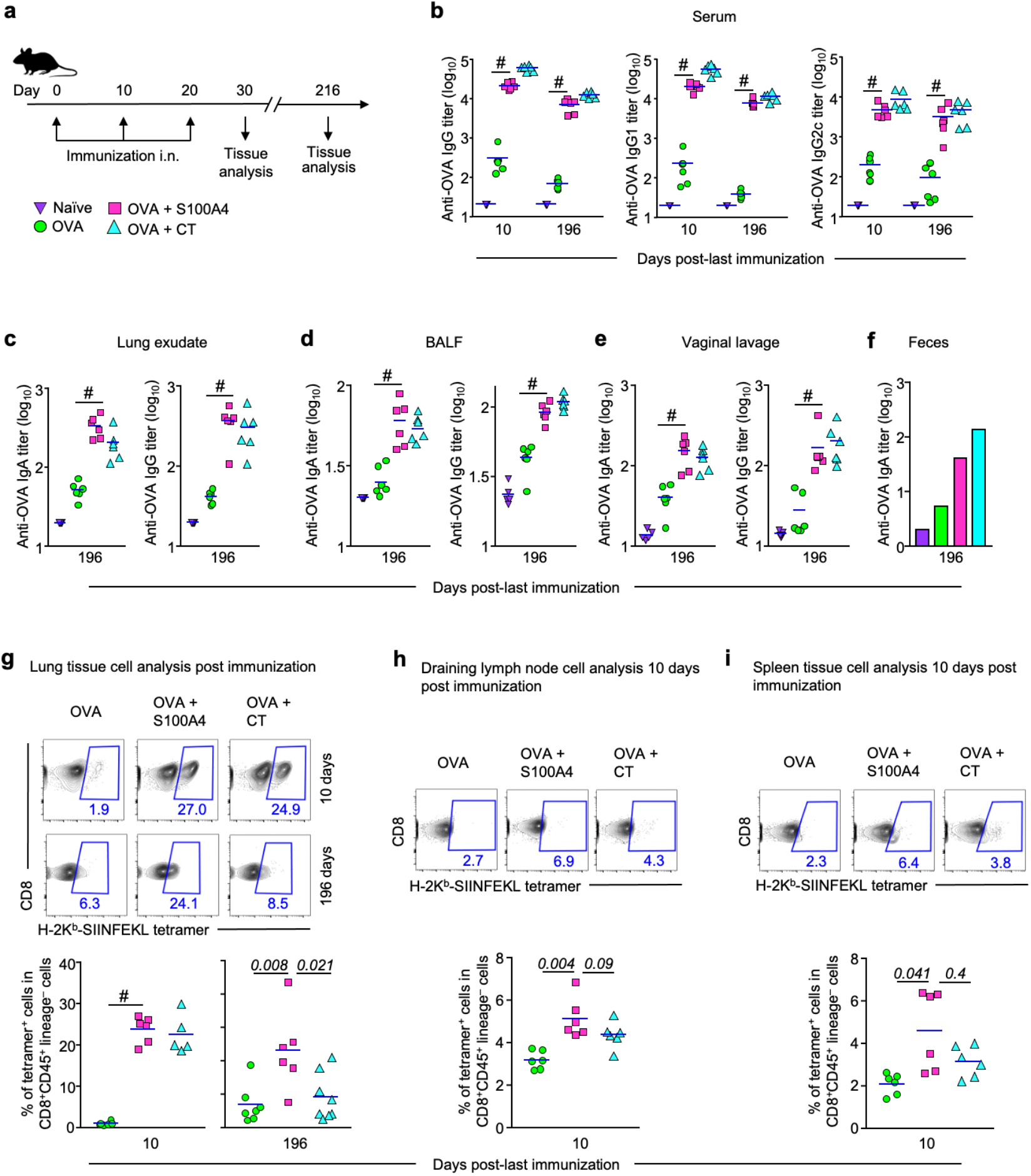
Intranasal immunization adjuvanted with S100A4 provokes greater humoral and cellular immune responses. Mice were immunized three times intranasally (i.n.) with ovalbumin (OVA; 10 μg) alone, or admixed to S100A4 (10 μg) or cholera toxin (CT; 1 μg) at a 10-day interval. Unmanipulated *naïve* mice were included for baseline control. Mice were euthanized 10 days or 196 days after the last immunization for collection of various types of samples. **a** The immunization and tissue collection procedures are indicated using a flowchart. **b-f** Various classes of OVA-specific antibody levels in serum **(b)**, lung exudates **(c)**, broncho-alveolar lavage fluid (BALF) **(d)**, vaginal lavage **(e)** and pooled feces **(f)** were analyzed by ELISA. **g-i** Single-cell suspensions were prepared from the lung tissue **(g)**, lymph nodes **(h)**, and spleen (i). Percentages of CD8 T cells that recognize H-2K^b^/OVA (SIINFEKL) tetramer were revealed by flow cytometry. Contour plots indicating a representative mouse from each treatment (upper panel) and pooled data from all the mice (lower panel) are shown. Numbers adjacent to outlined areas indicate percent cells in each gate. Each dot represents data from an individual mouse and blue lines indicate the average values. *#P* = 0.0022 or the exact P-values (italic numbers) are indicated by Mann-Whitney *U* test Except for the long-term observation experiment, the other measurements represent at least three separate experiments. Source data are available for this figure.

We also determined the accumulation of OVA_257-264_ peptide (SIINFEKL)-specific CD8 T cells in various tissues using the H-2K^b^-SIINFEKL tetramer analyzed by flow cytometry (Supplementary Fig. 2). Co-administration with S100A4 promoted the expansion of CD8 T cells that recognized the MHC class I-associated SIINFEKL peptide in the lung (Fig. 1g), lymph node (Fig. 1h), and spleen (Fig. 1i) at 10 days after the last immunization. A trend of enhanced expansion of these T cells in the lymph nodes and spleen was observed when comparing the effect of S100A4 with CT (Fig. 1h and i). For SIINFEKL peptide-specific T cell expansion in the lung, we also measured at 196 days after the last immunization, and S100A4-adjuvanted immunization maintained the response better than CT (Fig. 1g). These findings suggest that S100A4 could be helpful in driving cytotoxic T cell responses, in addition to stimulating humoral immunity, and may exert a critical role in maintaining protective immunity against viral infections.

To examine if the mucosal adjuvant activity of S100A4 on the immune response to the model antigen OVA would extend to a clinically relevant vaccine antigen, such as the spike protein of SARS-CoV-2 (“Spike”), we immunized mice with Spike alone or together with either S100A4 or CT as adjuvants using a similar intranasal immunization regimen as with OVA (Fig. 2a). Again, co-immunization with S100A4 resulted in substantially increased levels of Spike-specific total IgG and IgG1 antibodies in serum (Fig. 2b), as well as Spike-specific mucosal IgA responses in BALF (Fig. 2c), lung parenchyma (Fig. 2d), nasal mucosa (Fig. 2e), and eye mucosa (Fig. 2f). It has recently been reported that respiratory tract mucosal IgA exerts major neutralizing activity against SARS-CoV-2 *(Sterlin et al., 2020, Wang et al., 2020)*.

**Figure 2.**
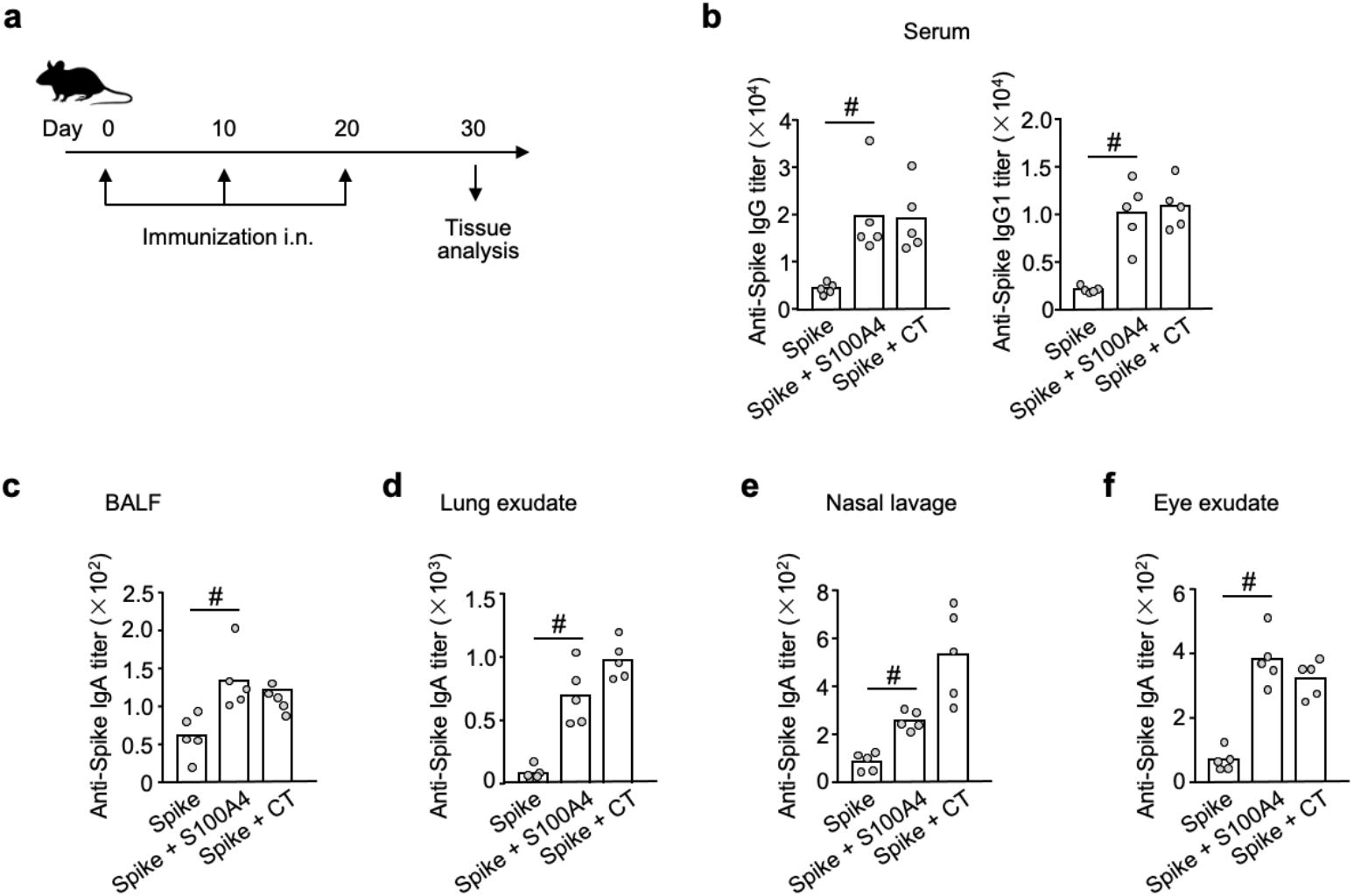
S100A4 promotes spike protein-specific antibody responses after intranasal immunization with the recombinant spike protein of SARS-CoV-2. Mice received three intranasal immunizations (i.n.) at an interval of 10 days. For each immunization, mice were intranasally administered with 5 μg spike protein alone or admixed with 10 μg S 100A4 or 1 μg cholera toxin (CT). Samples were collected 10 days after the last immunization **(a)**. Levels of anti-spike protein lgG and lgG1 in serum **(b)** as well as anti-spike protein lgA in broncho-alveolar lavage fluid (BALF) **(c)**, lung exudates **(d)**, nasal lavage **(e)**, and eye exudates **(f)** were measured by ELISA. Each dot represents measurement from an individual mouse and columns indicate the average values. *#P* = 0.0079 by Mann-Whitney *U* test. Shown is one of two similar experiments. Source data are available for this figure.

### S100A4 extends nasal residence time and promotes tissue transport of antigens

Nasal mucociliary clearance is a mechanism that limits the nasal residence time of vaccine antigens delivered intranasally *(Gizurarson, 2015)*, thus compromising mucosal antigen uptake. Efficient cell-mediated antigen transport to tissues and secondary lymphoid organs is critical to achieving vaccine efficacy *(Irvine et al., 2020)*. We performed *in vivo* real-time fluorescence optical imaging to monitor the kinetic biodistribution of intranasally administered fluorescently labeled OVA, either in the absence or presence of S100A4. Ventral and dorsal images of live mice were captured at various time points up until six hours after the administration of Alexa Fluor 647-conjugated OVA (OVA-AF647). Shortly after the intranasal administration of OVA-AF647, identical ventral view fluorescence signals were observed at the snouts of mice (5 min; Fig. 3a; ventral view). No fluorescence signals were detected from the dorsal side at this time point, probably due to blocking of the signal by the overlying turbinate bone. As can be clearly observed ventrally, the fluorescent signal persisted more robustly in the snouts of the mice that received the fluorescently labelled antigen together with S100A4, suggesting a longer nasal residence time of the antigen (Fig. 3a and b).

**Figure 3.**
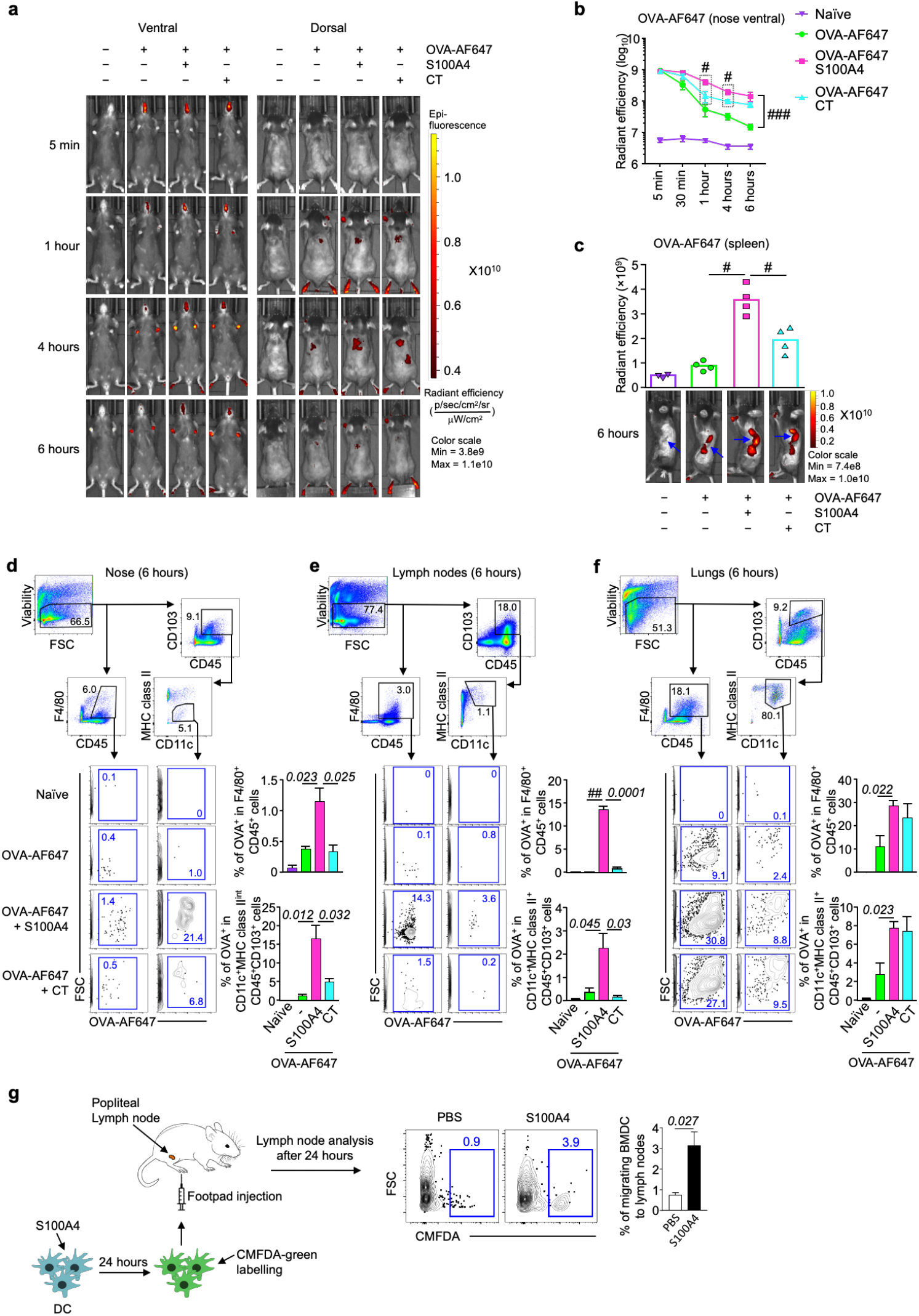
S100A4 promotes nasal antigen retention and antigen transport. **a-f** Mice received a single intranasal delivery of Alexa Fluor 647-conjugated ovalbumin (OVA-AF647; 10 μg) in the absence or presence of S100A4 (20 μg) or cholera toxin (CT; 1 μg). Unmanipulated naïve mice were included for background control. The ventral and dorsal images showing the biodistribution of OVA-AF647 in the mice were taken at various time points as indicated using an *in vivo* animal imaging system **(a)** and nasal retention of OVA- AF647 from the ventral view was quantified **(b)**. Mice were euthanized at six hours after antigen delivery and dorsal skin was cut open to expose the internal organs around the spleen site which was imaged **(c**, bottom; arrows indicate the spleen) and fluorescence accumulation in the spleen was quantified **(c**, top). Also at six hours after OVA administration, mouse nasal tissues **(d)**, cervical lymph nodes **(e)** and lungs **(f)** were harvested for flow cytometric analysis. Representative contour plots indicating the biodistribution of OVAAF647 in one representative mouse out of four in each treatment group are shown. **g** Bone marrow-derived dendric cells (BMDCs) were treated with or without S100A4 (1 μg/ml) overnight and stained with CellTracker™ (CMFDA-green). Mice were then injected with 2X 10^6^ BMDCs into the footpad and popliteal lymph nodes were harvested 24 hours later for flow cytometric analysis. Numbers in or adjacent to outlined areas indicate percent cells in each gate (d-f). Each dot represents measurement from an individual mouse (c). Data are expressed as mean + s.e.m. of four biological replicates **(b-g)**. *#P* = 0.0286 by Mann-Whitney *U* test (b-c). *###P* < 0.0001 by two-way ANOVA comparing the two groups for all the time points **(b).** *##P* < 0.0001 or the exact P-values (italic numbers) are indicated by unpaired *t*-test **(d-g).** Shown is one of two representative experiments. Source data are available for this figure.

Furthermore, co-administration with S100A4 enhanced the dorsal appearance of the fluorescence signals associated with tissues around the spleen, which started to appear at 30 min, peaked at around four hours, and declined at six hours post-antigen delivery (Fig. 3a; dorsal view). We then euthanized the mice at about six hours after antigen delivery and removed the skin at the left side of the abdominal cavity in order to get clearer views of the fluorescence deposition in relevant organs. The co-administration with S100A4 clearly promoted the accumulation of fluorescent OVA in the spleen, stomach, and kidney (Fig. 3c). In particular, the spleen demonstrated the most remarkable S100A4-driven antigen accumulation, as the mouse that received only fluorescent OVA showed merely the background fluorescence (Fig. 3c). For both the nasal retention and tissue accumulation of the fluorescent OVA, S100A4 demonstrated stronger effects than the CT used for comparison (Fig. 3a–c).

Next, we collected nasal tissues, cervical lymph nodes and lung tissues from each individual mouse at 6 hours after the delivery of fluorescent OVA and prepared single-cell suspensions to investigate, using flow cytometry, the distribution of the fluorescent OVA at the cellular level including the CD103^+^ migratory DCs *(Kedl et al., 2017)*. Co-administration with S100A4 profoundly promoted accumulation of migrating DC (CD45^+^CD103^+^CD11c^+^MHC II^high/int^)- and macrophage (CD45^+^F4/80^+^)-associated fluorescent OVA in the nasal tissue (Fig. 3d), cervical lymph nodes (Fig. 3e), and lung tissue (Fig. 3f). Of note, the antigen-associated DCs identified in the nasal tissue had a lower expression level of MHC class II. In contrast, CT only increased the cell-associated OVA accumulation in the lungs (Fig. 3f), but not nose (Fig. 3d) and lymph nodes (Fig. 3e) at six hours after antigen delivery. At 12 hours, potent effects of S100A4 on the accumulation of antigen-loaded antigen-presenting cells in the lungs could be revealed, while CT showed at least a trend of weaker effects at this time point compared with S100A4 (Supplementary Fig. 3). Efficient uptake and transport of vaccine antigens have been described as a critical quality for assessing the adjuvant activity *(Liang et al., 2017)*.

Using *in vitro* cultured bone marrow-derived DC (BMDC), we failed to observe any augmentation in the uptake of fluorescent OVA after treatment with either S100A4 or CT (Supplementary Fig. 4a). Similarly, uptake of fluorescent OVA by fresh *ex vivo* cells from the nasal tissue, cervical lymph nodes, and spleen was not enhanced by S100A4 or CT (Supplementary Fig. 4b). Thus, it is likely S100A4 promoted the migration of antigen-presenting DCs from the nasal mucosa but did not promote antigen uptake itself. Our *in vivo* imaging data showing S100A4-promoted antigen accumulation in the nasal tissue, cervical lymph nodes and lung tissues after nasal delivery of the antigen (Fig. 3d–f) support a stimulatory effect of S100A4 in promoting the migration of antigen-presenting cells including DCs and macrophages that had captured the antigen after mucosal delivery. To fundamentally confirm the effect of S100A4 on DC migration, we designed an adoptive transfer assay to unambiguously demonstrate more efficient DC migration driven by S100A4. S100A4-primed and control BMDCs were labelled with a green fluorescent dye for easy tracking. Following injection of the fluorescent DCs into the mouse footpad, substantially increased numbers of fluorescent DCs that had been previously treated with S100A4 arrived in the draining lymph nodes compared with the control DC (Fig. 3g).

### S100A4 effectively induces germinal center responses after intranasal immunization

The development of germinal centers is a recognized mechanism for the generation of high-quality antibodies after immunization *(Aradottir Pind et al., 2019, Lycke, 2010)*. We harvested mouse spleens and cervical lymph nodes after intranasal immunization for flow cytometric analysis of the impact of S100A4 on immune cells bearing the germinal center features using the gating strategy defined in Supplementary Fig. 5. Frequencies of activated marginal zone B cells (CD69^+^CD38^+^B220^+^) *(Oliver et al., 1997)* were increased in both the spleen and cervical lymph nodes of mice that received the antigen together with S100A4 (Fig. 4a). In both compartments, S100A4 also expanded germinal center B cells (GL7^+^FAS^+^CD38^−^B220^+^) (Fig. 4b) and T follicular helper cells (PD-1^+^CXCR5^+^CD4^+^Foxp3^−^), a cell type which is critical in the formation of germinal centers (Fig. 4c). The immune enhancing effects of S100A4 were similar to those of CT (Fig. 4 a– c). These results were confirmed using immune staining and confocal microscopy to demonstrate the typical germinal center morphology. Mouse spleens were first cryosectioned, followed by immunofluorescent staining for the expression of B220, GL7, as well as the cell proliferation marker Ki-67, as previously reported *(Sun et al., 2017)*. Immunization with S100A4 strongly promoted the expansion of GL7^+^ and Ki-67^+^ B cells and morphological changes associated with germinal center formation (Fig. 4d).

**Figure 4.**
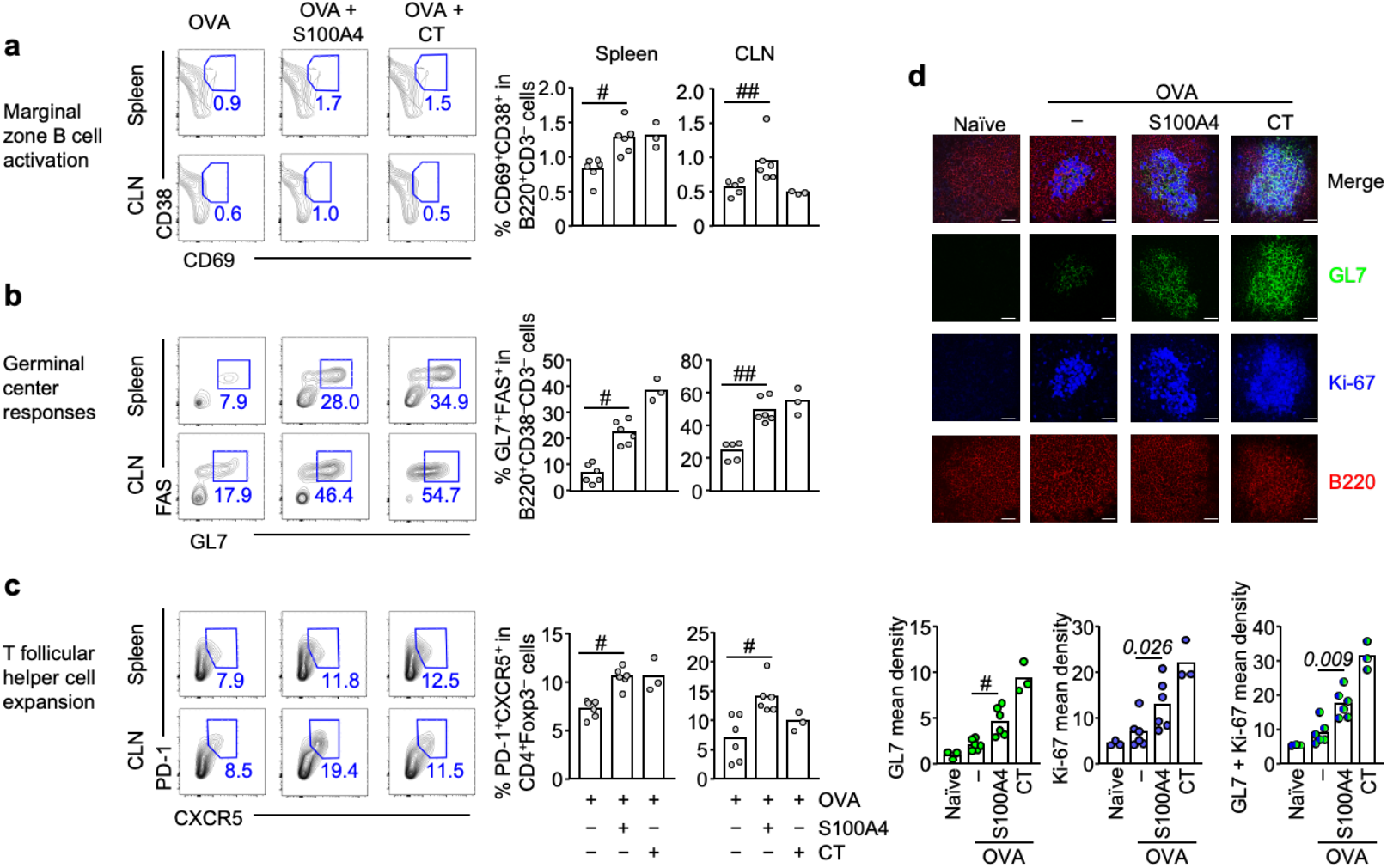
Intranasal immunization adjuvanted with S100A4 activates secondary lymphoid organ cells and promotes germinal center formation. Mice received three intranasal immunizations as explained in Supplementary Figure 1A. Spleens and cervical lymph nodes (CLN) were collected three days after the last immunization. **a-c** Single cell preparation was analyzed by flow cytometry. Frequencies of CD69^+^CD38^+^ activated marginal zone B cells **(a)**, GL 7^+^FAS^+^CD38^−^B cells that characterize germinal center formation **(b**), and **PD-1** ^+^CXCR5^+^Foxp3^−^T follicular helper cells **(c)** were analyzed. Each contour plot indicates data from one representative mouse and numbers in or adjacent to outlined areas indicate percent cells in each gate (left panels). The bars show the pooled data (right panels). **d** Spleen sections were analyzed by immunohistochemistry to visualize germinal center morphology. Red, B220; green, GL 7; blue, Ki-67. The top panels of the microscopic pictures show merged images. The bottom bar figure panels show the quantification of staining intensities. Scale bar, 25 μm. Each dot represents data from an individual mouse and columns indicate the average values. *#P* = 0.0022; *##P* < 0.0043 or the exact P-values (italic numbers) are indicated by Mann-Whitney *U* test. Shown is one of two representative experiments. Source data are available for this figure.

### S100A4-adjuvanted intranasal immunization promotes lipid accumulation in the spleen

It has recently been reported that germinal center B cells require active oxidation of fatty acids instead of glycolysis to meet the energetic challenge of rapid cell proliferation *(Weisel et al., 2020)*. In order for efficient germinal center development to take place, hosts need to mobilize lipid transport to the germinal centers. Therefore, we examined the accumulation of lipids using MALDI-TOF MS after S100A4-adjuvanted immunization. A variety of lipids, with glycerolipids and glycerophospholipids being the dominant forms, were identified in the spleens (Supplementary Fig. 6). A shotgun lipidomics approach was used to analyze all lipid classes together. The abundance of total lipids was increased in the spleen after co-immunization with S100A4 (Fig. 5a). The lipid intensities were quantified and expressed using an intensity box plot which displayed a remarkable increase of lipids in spleens from S100A4-adjuvanted mice (Fig. 5b). Principal component analysis revealed a complete separation of S100A4-adjuvanted immunization from both the OVA only immunization and the naïve mouse controls (Fig. 5c). A similar and more intense lipid accumulation response than that induced by S100A4 was seen in mice given CT together with OVA (Fig. 5).

**Figure 5.**
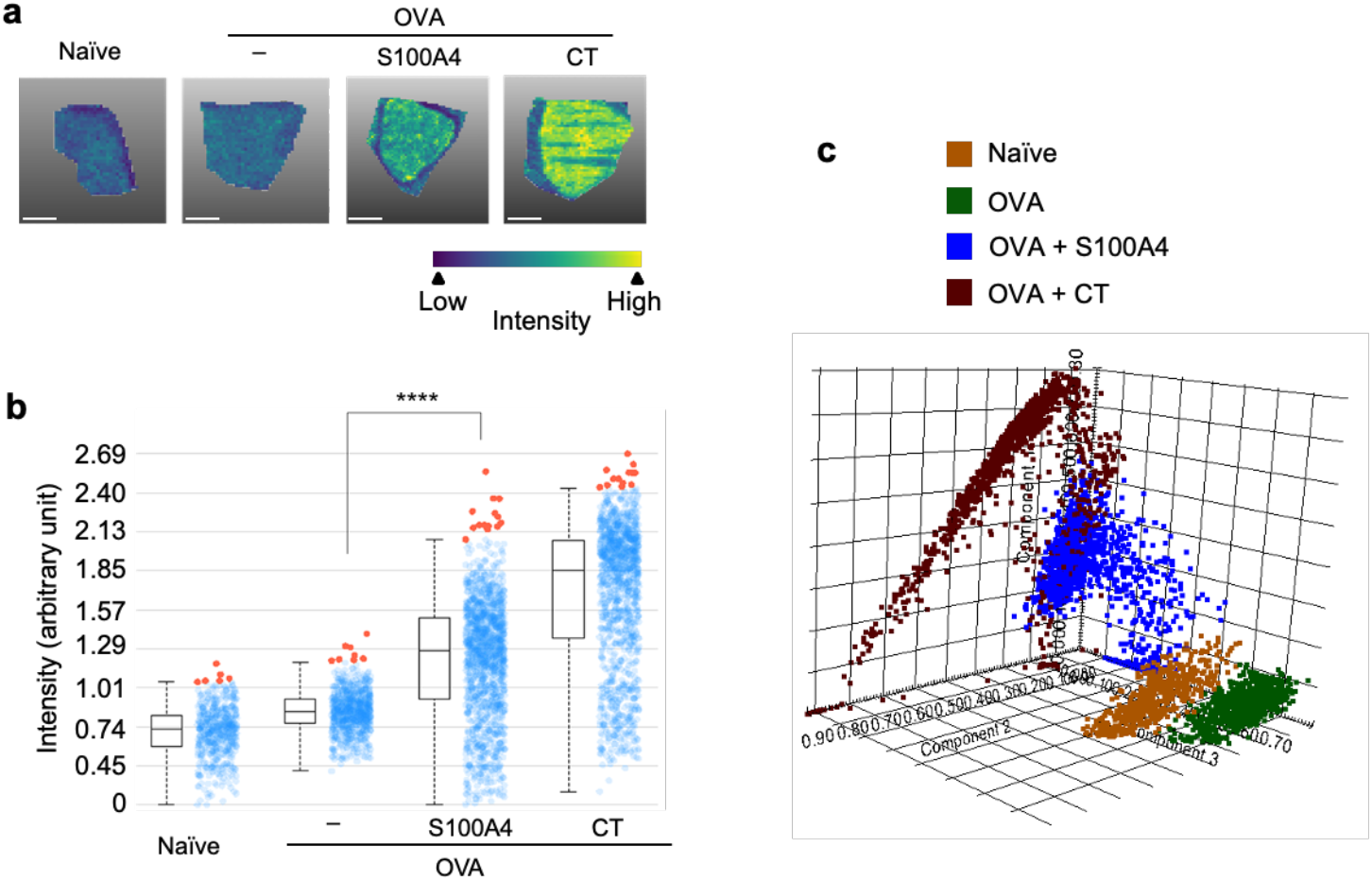
Intranasal immunization adjuvanted with S100A4 promotes lipid accumulation in the spleen. Mice received three intranasal immunizations as explained in Supplementary Figure 1A. Spleens were collected three days after the last immunization and sectioned for MALDI-TOF analysis. **a** The intensity and distribution of lipids within a range of mass-to-charge (m/z) ratio (754.8 ± 314.3 Da) are shown as MALDI MSI ion pseudo-colour images. Scale bar, 6 mm. **b** Expression of overall levels of the identified lipids were quantified by intensity box plots. The intensity of a mass peak from each mass spectrum is depicted by a spot within the plot, and the outliers are coloured red. **c** Principal component (PC) analysis of MALDI MSI data using the score plot of the three first principal components (PC1-PC3) displays various lipid mass spectra highlighting different lipid signatures between various treatment groups. Results show one representative mouse **(a)** or represent three to six mice **(b, c)** in each treatment group. *\*\*\*\*P* < 0.0001 by Kruskal-Wallis test **(b)**. OVA, ovalbumin.

As our germinal center activity investigation using various approaches was based on the same cohort of mice, we compared the lipid and GL7 germinal center marker measurements in individual mice. Good correlations were established between the expression of GL7 using flow cytometry and fluorescence microscopy and the lipid abundance determined using MALDI-TOF (Supplementary Fig. 7), supporting the usefulness of lipid abundance as a surrogate marker for the germinal center response.

### Extracellular S100A4 potentiates DC activation

DCs are critical antigen-presenting cells that bridge innate and adaptive immune responses and, as such, are rational targets for the choice of effective adjuvants *(Tesfaye et al., 2019)*. We have previously demonstrated that adoptive engraftment of wild-type DCs could restore the defective adaptive immune responses of S100A4-deficient mice *(Sun et al., 2017)*. Here we examined the regulatory effect of exogenous S100A4 on cultured DCs. Our data clearly demonstrates that mRNA expression of a group of cytokines and molecules with immune regulatory activities was upregulated in BMDCs following incubation with S100A4 (Fig. 6a). Specifically, the transcript levels of a number of cytokines that are critical to adaptive immunity, including IL-1β, IL-2, IL-6, and IL-10, were augmented after treatment with S100A4, as were co-stimulatory molecules, including CD80, CD86, and CD40. Furthermore, at the protein level, S100A4 was found to potently activate BMDCs by augmenting the production of these costimulatory molecules, as well as IL-1β and IL-6 (Fig. 6b).

**Figure 6.**
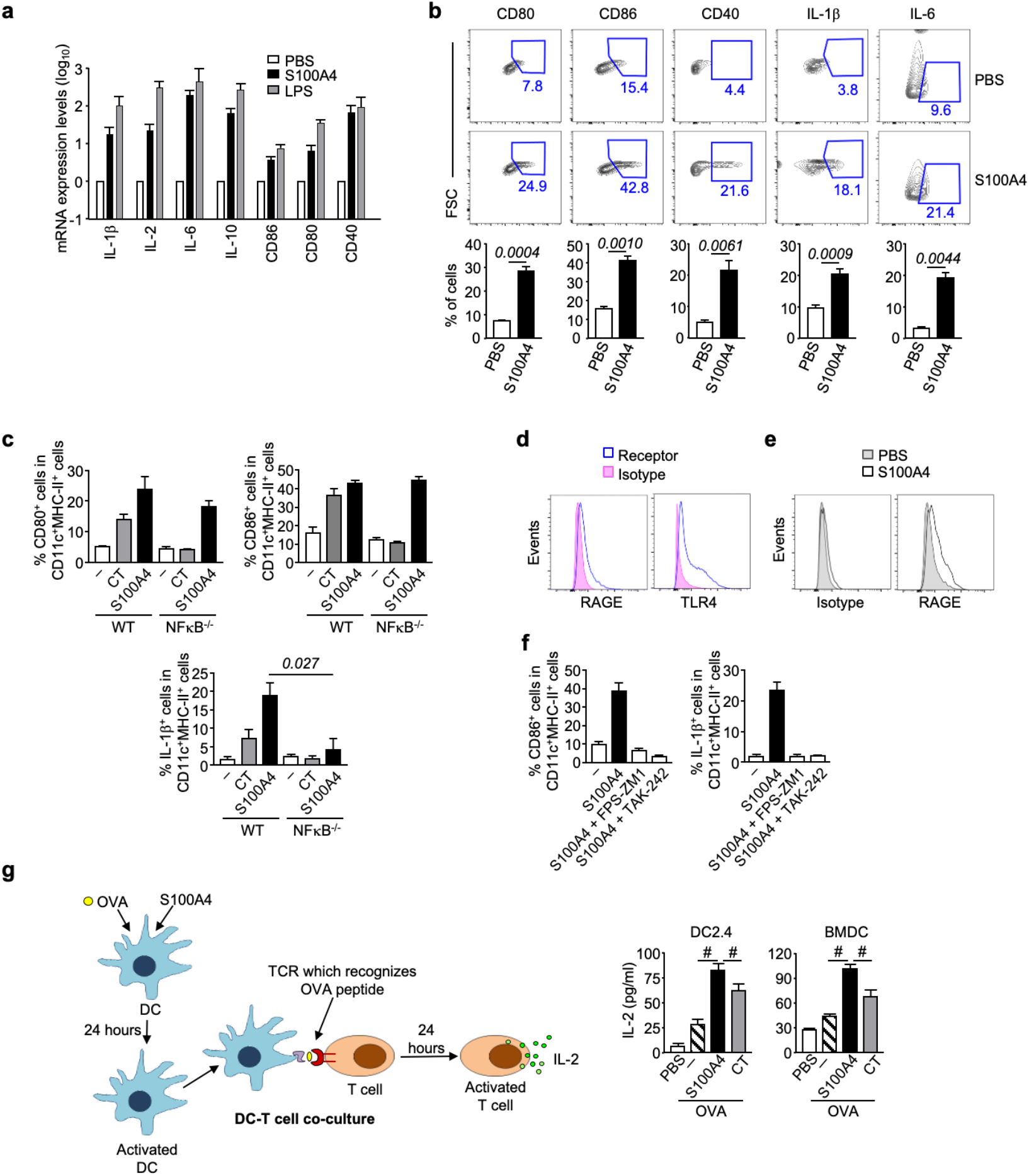
5100A4 activates dendritic cells. **a, b** Bone marrow-derived dendritic cells (BMDCs) were incubated with or without 1 μg/ml S100A4 for three hours **(a)** or overnight **(b)**. Some cells were stimulated with 1 μg/ml LPS as a positive control. Expression of a number of molecules important for DC activation as indicated was analyzed at the mRNA level by quantitative reverse transcription PCR **(a)** or at the protein level by flow cytometry **(b;** the lower panel shows the quantification). **c** BMDCs were cultured from wild-type or NFKB-deficient mice followed by incubation overnight with or without S100A4 or cholera toxin (CT; 0.1 μg/ml). Cell surface expression of CD80 and CD86, and intracellular production of IL-1 ~ were measured by flow cytometry. **d** BMDCs were incubated with anti-RAGE (left panel) or anti-TLR4 (right panel), or with isotype control antibodies, followed by flow cytometric analysis. **e** BMDCs were incubated with or without S100A4 and their surface expression of RAGE was determined as explained in **(d). f** BMDCs were incubated overnight with S100A4 or S100A4 pre-mixed with FPS-ZM1 or TAK-242, inhibitors of RAGE and TLR4, respectively. Cell surface expression of CD86 (left panel) and intracellular production of IL-1~ (right panel) were measured by flow cytometry as explained in **(b). g** BMDCs or a DC cell line (DC2.4) were treated with OVA (1 mg/ml) mixed with or without S100A4 (1 μg/ml) or CT (1 μg/ml) overnight before co-culturing with RF33.70 hybridoma (T cells with T cell receptor specific for the OVA_257.264_ peptide) at a 1 :1 (BMDC to RF33.70) or 1 :10 (DC2.4 to RF33.70) ratio overnight. T cell activation was quantified by measuring IL-2 release using ELISA, with a schematic diagram showing the experimental design. Expression was normalized using GAPDH as the calibrator gene **(a)**. Representative data out of at least three separate BMDC cultures are shown **(d, e)**, or data are expressed as mean+ s.e.m. of three to four individual cell cultures **(a, b, c, f, g)**. *#P* = 0.0022 or the exact P-value (italic number) is indicated by Mann-Whitney *U* test. The exact P-values (italic numbers) are indicated by unpaired *t*-test **(b)**. Source data are available for this figure.

We have previously shown a dependence on NFκB signaling as a critical DC activation pathway used by CT to exert mucosal adjuvant activity *(Terrinoni et al., 2019)*. Here, we can also demonstrate that S100A4-mediated DC activation is at least partially dependent on NFκB signaling. Thus, S100A4-induced production of IL-1β, a critical molecule in the pathway of adjuvanticity, was substantially suppressed in NFκB^−/−^ BMDCs, although the increased expression of CD80 and CD86 by S100A4 remained intact in NFκB^−/−^ cells (Fig. 6c). The latter finding may suggest that S100A4 could have a broader spectrum of signaling pathways than CT, which failed to increase both costimulatory molecule expression and IL-1β production in NFκB^−/−^ BMDCs (Fig. 6c). Extracellular S100A4 has been shown to activate NFκB in various types of cells *(Boye et al., 2008, Kim et al., 2017, Schmidt-Hansen et al., 2004)*. Therefore, it is not surprising that S100A4, as shown here, also stimulates DCs through the NFκB pathway.

It is reported that S100A4 triggers intracellular signaling pathways through RAGE and TLR4 *(Bjork et al., 2013)*. Indeed, BMDCs expressed both RAGE and TLR4 (Fig. 6d), and we found that RAGE expression was upregulated by exogenous S100A4 (Fig. 6e). We then tested whether S100A4-mediated DC activation would require either or both of these two receptors. To this end, BMDCs were incubated with S100A4 either in the presence or absence of FPS-ZM1, an inhibitor of RAGE, or TAK-242, an inhibitor of TLR4, and the effects on CD86 expression as well as production of IL-1β were measured. The results showed that S100A4-induced upregulation of both of these activation markers was almost completely abrogated by blocking either RAGE or TLR4 (Fig. 6f), suggesting the requirement of both receptors for S100A4-induced DC activation.

### Extracellular S100A4 enhances MHC class I antigenic peptide cross-presentation by DC and activation of cytotoxic T cells

We have demonstrated the potent effect of S100A4 in stimulating cytotoxic CD8 T cells against a co-administered antigen (Fig. 1g–i). Next, we wished to also confirm that S100A4 could enhance MHC class I-associated antigenic peptide cross-presentation on DCs, a critical step in the activation of cytotoxic T cell responses after immunization with protein-based subunit vaccines *(Ho et al., 2018)*. To this end, we incubated OVA-pulsed DCs either in the presence or absence of S100A4, followed by co-culturing the DCs with RF33.70 hybridoma T cells that had T cell receptors specific for OVA_257-264_ presented by H2K^b^. S100A4-pulsed DCs substantially augmented the activation of T cells as measured by IL-2 production (Fig. 6g). S100A4 demonstrated a more potent effect compared with CT (Fig. 6g).

### The adjuvanticity of S100A4 is not affected by contaminating lipopolysaccharide (LPS)

We performed both *in vivo* immunization and *in vitro* DC activation assays to rule out the possible confounding effect of a residual low amount of endotoxin present in the recombinant S100A4 product (10 pg LPS per μg S100A4, as measured using a Limulus amebocyte lysate assay). *In vivo*, mice were given three intranasal immunizations at ten-day intervals with OVA in the presence of various doses of S100A4 or various amounts of LPS corresponding to the residual LPS amount in a given S100A4 dose. Ten days after the last immunization, blood was collected and OVA-specific total IgG antibody production in serum was measured. The tested range of LPS amounts failed to promote OVA-specific IgG production, supporting that the enhancement of this response by S100A4 is a true adjuvant effect of S100A4 but not the result of contaminating LPS (Supplementary Fig. 8a). On the other hand, as a positive control, LPS at 1 μg (5000-fold of the highest contamination level) effectively promoted the anti-OVA response (Supplementary Fig. 8a).

In the *in vitro* experiments, we incubated BMDCs with LPS at 10 pg/ml, i.e. the estimated contamination level of LPS in 1 μg/ml S100A4, which was tested in parallel, and observed no enhancement in the expression of the genes under study (Supplementary Fig. 8b). In contrast, a 100,000-fold higher level of LPS (1 μg/ml) demonstrated potent stimulatory effects (Supplementary Fig. 8b). Further support for the S100A4 protein, but not the residual endotoxin contamination, being the entity responsible for DC activation came from studies showing that the strong promotion of CD80, CD86 and IL-1β in BMDCs by the recombinant S100A4 was largely suppressed by a neutralizing antibody against S100A4 (Supplementary Fig. 8c).

### S100A4 does not induce humoral immune responses against itself after intranasal immunization

In vaccination practice, an ideal adjuvant should be able to support different vaccine antigens in multiple administrations, so that the adjuvant can be formulated in different vaccine preparations. Therefore, immune responses against the adjuvant itself should preferably be avoided. To assess the quality of S100A4 in this respect, we repeated our basic intranasal immunization model using OVA as an experimental antigen admixed with either S100A4 or CT as an adjuvant (Supplementary Fig. 9a). Consistent with the previous findings (e.g., Supplementary Fig. 1b and Fig. 1b), we again showed that both S100A4 and CT were able to remarkably augment anti-OVA IgG levels (Supplementary Fig. 9b and c). Interestingly, while CT, as a foreign protein, stimulated substantial levels of anti-CT IgG antibody responses (Supplementary Fig. 9c), S100A4 failed to stimulate anti-S100A4 IgG antibody production above the level seen in naïve mice (Supplementary Fig. 9b). This could be due to the fact that S100A4 is an internal protein subject to immune tolerance. The data suggests that S100A4 could lead to improved tolerability over non-host proteins (e.g., CT) in repeated use as a vaccine adjuvant.

## Discussion

The search for clinically applicable mucosal adjuvants still remains a challenging endeavor. By virtue of their strong cAMP mobilization capacity, both CT and heat-labile enterotoxin (LT) from *Escherichia coli* have been used as experimental mucosal adjuvants. Although for safety reasons, these toxins are not suitable as mucosal adjuvants in humans *(Lycke, 2012)*, CT has been extensively studied and is considered the “gold standard” experimental mucosal adjuvant *(Sanchez & Holmgren, 2011)*. Therefore, CT is commonly used as a benchmark for comparing the adjuvant effects of other candidate mucosal adjuvants. Various approaches have been attempted to modify these native bacterial toxins to derive mutants that can be used clinically *(Lycke & Lebrero-Fernandez, 2018)*. Yet, toxin-independent approaches are also worth exploring. In this study, we present experimental evidence that nasal immunization with protein antigens adjuvanted with S100A4 induced adaptive immune responses comparable, or superior, in magnitude to those achieved using CT as an adjuvant. S100A4 augmented antigen-specific immune responses, both systemically and at mucosal surfaces, to intranasally co-administered antigens without any adverse effects, as shown using both OVA as a model antigen and the SARS-CoV-2 spike protein as a clinically applicable vaccine antigen. Furthermore, S100A4 potently induced the expansion of antigen-specific CD8 T cells, which could persist in lung tissues for at least six and a half months after immunization, similar to the persistence of the high titers of antigen-specific antibodies in various compartments. The fact that exogenous S100A4 was demonstrated to effectively prolong mucosal antigen retention, mobilize DC and macrophage migration, promote germinal center responses, and activate cytotoxic T cells, supports the proposed usefulness of this molecule as a mucosal adjuvant.

S100A4 is a member of the damage-associated molecular pattern (DAMP) family *(Bertheloot & Latz, 2017)*. This heterogeneous family is composed of proteins, lipids, nucleic acids, and small-molecule compounds that are normally sequestered inside cells but are released upon stress stimulation, e.g., tissue injury *(Venereau et al., 2015)*. DAMP molecules have been implicated in the activation of inflammasomes *(Lage et al., 2014)*, a process used by many adjuvants *(Ivanov et al., 2020)*. We have previously demonstrated that DCs from mice lacking S100A4 have compromised inflammasome-associated caspase-1 and IL-1β production upon treatment with CT *(Sun et al., 2017)*. In the present study, we show that extracellular S100A4 can promote not only the upregulation of critical costimulatory molecules in DCs, including CD80, CD86, and CD40, but also the production of key cytokines, including IL-1β, for the induction and maintenance of adaptive immunity. The generation of IL-1β by DCs and other APCs as a consequence of NFκB activation and an inflammasome response has been found to be a critical step in the adjuvant function of CT *(Terrinoni et al., 2019)*. Taken together, our work suggests not only an endogenous effect of S100A4 in mucosal immunization *(Sun et al., 2017)*, but also that, as a secreted protein *(Schmidt-Hansen et al., 2004)*, S100A4 may have a paracrine or endocrine function on cells including APCs, which is likely further enhanced by the exogenous S100A4 added as an adjuvant.

In addition to the possible induction of mucosal tolerance, poor uptake and rapid clearance by mucociliary movement of the antigen at the nasal mucosal surface have been described as hurdles for developing effective nasal immunization strategies *(Gizurarson, 2015)*. Various types of mucoadhesive agents have been co-administered with the antigens to improve nasal retention and antigen uptake *(Jabbal-Gill, 2010)*. Chitosan, as a biocompatible and biodegradable natural polymer, has been extensively investigated as a safe and effective mucoadhesive for improving nasal vaccine delivery *(Kang et al., 2009)*. Interestingly, as shown here, S100A4, in addition to its potent immune stimulatory effect, remarkably prolonged the nasal residence time of co-administered fluorescent OVA. This effect is likely due to the calcium-binding activity of S100A4, since free calcium is critical for maintaining the ciliary beating frequency *(Schmid & Salathe, 2011)*, and restricted ciliary beating may lead to a prolonged antigen residence time. In addition to prolonging the mucosal antigen retention time, our data also suggests that S100A4 can promote DC migration from the nasal mucosa. The latter effect may depend on the ability of S100A4 to both increase and engage RAGE on DCs, insofar as it has been previously observed that DCs depend on RAGE for their *in vivo* homing to lymph nodes *(Manfredi et al., 2008)*.

We further demonstrated that nasal co-administration of an antigen with S100A4 strongly promoted germinal center formation in both draining lymph nodes and the spleen. It is in the germinal center that high-affinity B cells and long-lived plasma cells are formed *(Shlomchik & Weisel, 2019, Suan et al., 2017)*, and their clonal expansion involves vigorous energy-demanding cell proliferation. In turn, this requires an adequate energy supply through activation of relevant metabolic pathways *(Jung et al., 2019, Waters et al., 2018)*. Weisel et al. recently demonstrated that proliferating germinal center B cells use fatty acids for energy supply instead of aerobic glycolysis to meet their metabolic requirements *(Weisel et al., 2020)*. This finding indicates that a successful germinal center reaction requires a substantial input of fatty acids which can be generated by lipid metabolism. Prompted by this discovery, we undertook to quantitatively assess lipid accumulation in the spleen in response to adjuvant-enhanced germinal center formation using MALDI-TOF MS. Our results demonstrate that the combined intensity of various lipid species in immunized mice adjuvanted with S100A4 was dramatically increased. Importantly, lipid accumulation in the spleen, as measured by MALDI-TOF, correlated closely with the levels of the established germinal center marker antigen GL7, measured using conventional techniques including flow cytometry and fluorescence microscopy. Taken together, our data suggests that lipid measurement by MALDI-TOF can provide a surrogate marker of germinal center activity. Being a label-free technology, MALDI-TOF could be a valuable tool for quantification of germinal center activity, not least in non-rodent animal models, in which the availability of commercial antibodies for specific germinal center immunostaining is lacking.

S100A4 is expressed in many normal cells, including fibroblasts, endothelial cells, smooth muscle cells, and leukocytes *(Boomershine et al., 2009, Li et al., 2010, Mishra et al., 2012, Strutz et al., 1995)*. Furthermore, S100A4 has been found to be critical in maintaining the normal biological functions of various types of cells through regulating cell growth, survival, differentiation, and motility *(Donato et al., 2013)*. We note here that clinically-oriented research on S100A4 has largely focused on its cancer metastasis-promoting properties *(Boye & Maelandsmo, 2010, Mishra et al., 2012)*. However, this does not preclude its applicability as an adjuvant in vaccination, as there is no evidence that exogenous S100A4 can induce cancer metastasis, and, on the contrary, S100A4 is also found to exert anticancer activity *(Dukhanina et al., 2018)*. Importantly, we did not observe any obvious side effects either shortly after or more than six months after nasal administration with S100A4. Therefore, we believe that S100A4, as a human endogenous protein, may have an advantageous safety profile compared with microbial pathogen-associated molecular pattern (PAMP) molecules, which have been extensively tested for possible modification as a mucosal adjuvant *(Demento et al., 2011)*. Notwithstanding this assumption, rigorous safety evaluations must be undertaken before S100A4 can be used clinically as a vaccine adjuvant.

## Materials and Methods

### Experimental animals

Wild-type C57BL/6 mice were purchased from B&K Universal AB, Stockholm, Sweden, or bred in-house at the Centralized Animal Facilities at the Hong Kong Polytechnic University. NFκB p50^−/−^ mice on a C57BL/6 background were purchased from the Jackson Laboratory. Female mice were six to eight weeks old at the start of all experiments. The studies were approved by the Ethical Committee for Laboratory Animals in Gothenburg, Sweden, and the Animal Subjects Ethics Sub-Committee of the Research Committee at the Hong Kong Polytechnic University.

### Immunization and collection of specimens

OVA (grade V) was purchased from Sigma-Aldrich. Recombinant mouse S100A4 was purchased from Gentaur Molecular Products. The residual amount of endotoxin in the recombinant S100A4 preparation was examined using a Pierce™ Chromogenic Endotoxin Quant Kit (Thermo Fisher Scientific). Mice were anesthetized using isoflurane prior to immunization. For intranasal immunization, mice received a total of 20 μl (10 μl in each nostrial) phosphate-buffered saline (PBS) containing 10 μg OVA or 5 μg of the spike protein of SARS-CoV-2 (the receptor-binding domain; recombinant product from Sino Biological) mixed with or without 10 μg or 20 μg S100A4 as indicated. In some experiments, LPS (various amounts as indicated) or CT (1 μg) were used as control adjuvants. Three different immunization models were constructed based on when samples were harvested for analysis (3, 10, or 196 days). For all three models, three intranasal immunizations were performed, with samples being harvested after the third (final) immunization. Samples included serum, lungs, BALF, nasal lavage, vaginal lavage, eye exudate, fecal extracts, cervical lymph nodes, and spleens.

### Antigen-specific antibody measurement

Enzyme-linked immunosorbent assay (ELISA) was employed to measure the titers of antigen-specific antibodies. A 96-well ELISA plate (Thermo Fisher Scientific) was coated with 100 μl OVA (100 μg/ml), the spike protein (2 μg/ml), S100A4 (2 μg/ml), or CT (2 μg/ml), followed by an overnight incubation at 4°C. After washing and blocking, specimens from immunized or control mice were added after appropriate serial dilution followed by incubation at 37°C for two hours. After rigorous washing, the plate was incubated with goat anti-mouse secondary antibodies for IgG (Southern Biotech; 1030-05), IgG1 (Southern Biotech; 1070-05), IgG2c (Southern Biotech; 1079-05), or IgA (Southern Biotech; 1040-05) conjugated with horseradish peroxidase, followed by standard color development. Absorbance measurements and titer calculation were completed using a BMG SPECTROStar Nano microplate reader.

### Cell culture and treatment with S100A4

BMDCs were generated by culturing bone marrow cells in culture medium (RPMI-1640; Sigma-Aldrich) supplemented with 10% fetal bovine serum (Gibco), 1% L-glutamine, 1% gentamicin, and 50 μM 2-mercaptoethanol (all from Sigma-Aldrich) in the presence of 200 ng/ml recombinant Flt3L (PeproTech) for nine days. Cells were incubated *in vitro* with various stimuli, including CT (List Biological Laboratories) (0.1 μg/ml), recombinant S100A4 (1 μg/ml), or LPS (Sigma-Aldrich)(10 pg/ml or 1 μg/ml), for specified time periods at 37°C in 5% CO2, followed by washing and re-suspension in buffer for flow cytometric examination or RNA purification. Where indicated, an anti-S100A4 antibody (clone 6B12) *(Klingelhofer et al., 2012)* or isotype control was used to block functional S100A4 extracellularly. To investigate whether the surface expression of the receptors RAGE and TLR4 were required for S100A4 to activate DCs, BMDCs were incubated overnight with S100A4 or S100A4 pre-mixed with FPS-ZM1 (10 μM; Sigma-Aldrich) or TAK-242 (10 μM; Calbiochem), inhibitors of RAGE and TLR4, respectively. For the *in vitro* uptake of OVA and presentation of the peptide by MHC class I on DC, BMDCs were incubated with OVA-AF647 (Thermo Fisher Scientific; 1 μg/ml) in the presence or absence of S100A4 (1 μg/ml) or CT (0.1 μg/ml), followed by flow cytometric analysis of the uptake of fluorescent OVA and the presentation of the OVA_257-264_ (SIINFEKL) by mouse MHC-I (H2K^b^) using a PE-conjugated anti-H2K^b^-SIINFEKL antibody (Biolegend; clone 25-D1.16). To further confirm the role of S100A4 in promoting antigen cross-presentation, BMDC (1 × 10^5^ cells/well) or DC2.4 (a DC cell line; Sigma-Aldrich) (1 × 10^4^ cells/well) were treated with OVA (1 mg/ml) mixed with or without S100A4 (1 μg/ml) or CT (1 μg/ml) overnight. After washing, DCs were co-cultured with RF33.70 cells (1 × 10^5^ cells/ well), a T-T hybridoma with T cell receptor (TCR) specific for OVA_257-264_ (SIINFEKL) presented by H-2K^b^ (a gift from Dr KL Rock) *(Rock et al., 1990)* at 1:1 (BMDC to RF33.70) or 1:10 (DC2.4 to RF33.70) overnight. Release of IL-2 into supernatant was measured using a commercial ELISA kit (Sigma-Aldrich).

### Live mouse imaging

The dorsal skin of mice was shaved to prevent attenuation of the fluorescent signal. Mice were administered intranasally with 20 μl PBS containing 10 μg of OVA-AF647 alone or mixed with 20 μg of S100A4 or 1 μg CT. The fluorescence intensity was immediately measured using an IVIS Lumina Series III pre-clinical *in vivo* animal imaging system (Perkin Elmer), and the measurement was repeated at various time points for six hours. Results were analyzed using Living Image 4.7.3 (Perkin Elmer). Naïve mice were used to adjust settings for eliminating background fluorescence.

### *In vivo* migration assay

To investigate DC migration, BMDCs were treated with or without S100A4 (1 μg/ml) overnight and stained with CellTracker™ green (5-chloromethylfluorescein diacetate [CMFDA]; Thermo Fisher Scientific) according to the manufacturer’s instruction. 2 × 10^6^ stained cells were injected into the footpad of naïve 8-12 weeks old C57BL/6 mice and popliteal lymph nodes were harvested after 24 hours for flowcytometric analysis.

### Flow cytometric analysis

*In vitro* cultured cells or single cell suspensions from various types of tissues were prepared. For staining of surface markers, cells were incubated with fluorescent antibodies to mouse B220, CD3, CD4, CD11b, CD11c, CD38, CD69, CD80, CD86, CD40, GL7, FAS, CXCR5, MHC class II, Ter-119, GR-1, and CD19 and PD-1 (BD Biosciences or eBioscience). Dead cells were excluded by staining with BD Horizon™ Fixable Viability Stain 620. For measuring intracellular expression of IL-1β, TNF-α, or Foxp3, cells were first stained for relevant surface markers as described above, then fixed and permeabilized using the Intracellular Fixation and Permeabilization Buffer Set (BD Biosciences), followed by intracellular staining with antibodies against IL-1β (clone NJTEN3; eBioscience), TNF-α (clone TN3-19.12; eBioscience), or Foxp3 (clone FLK-16; Nordic Biosite). Cell surface expression of the receptors RAGE and TLR4 was determined by staining the cells with anti-RAGE and anti-TLR4 or their respective isotype control antibodies. BV421-conjugated H-2K^b^-SIINFEKL tetramer (MBL International) was used for the measurement of CD8 T cells that can recognize OVA_257-264_ peptide (SIINFEKL) presented by MHC class I. Cells were analyzed using an LSR II or FACSAria III flow cytometer (BD Biosciences). The data were analyzed using FlowJo software (Tree Star).

### Confocal microscopy

Mouse spleens were embedded in Cryomatrix™ embedding resin (Thermo Fisher Scientific). Frozen tissues were sectioned at 7 μm thick using CryoStar™ NX70 Cryostat (Thermo Fisher Scientific), then fixed in acetone, air-dried, and rinsed in PBS. Next, spleen cryosections were stained with anti-B220-biotin (BD Biosciences), GL7-Alexa Fluor 488 (eBioscience), and Ki-67-Horizon V450 (BD Biosciences), followed by incubation with Alexa Fluor 594-conjugated streptavidin (Invitrogen). Slides were mounted with ProLong™ gold antifade mounting media (Thermo Fisher Scientific). Confocal images were acquired using a TCS SPE confocal microscope (Leica) with a 60 × zoom. The data were analyzed using LAS X Leica software (Leica).

### MALDI-TOF MS analysis

Spleen tissues were immediately frozen in liquid nitrogen and stored at −70°C, followed by sectioning, as described above. Sections were embedded onto an indium tin oxide (ITO)-coated conductive glass slide and left to dry in a vacuum desiccator overnight. A layer of MALDI matrix [2,5-dihydroxybenzoic acid (30 mg/ml) dissolved in a mixture of 70% acetonitrile and 30% water containing 0.1% trifluoroacetic acid] was applied by aerosol spraying onto the dried slides. Matrix-coated slides were then mounted on the MALDI target adapter after wiping off the matrix from the outer regions of the slide using a tissue wetted with ethanol to ensure optimum electric contact with the adapter. Slides were then analyzed for lipids and fatty acids using an UltrafleXtreme MALDI-TOF mass spectrometer (Bruker). The results were processed using SCiLS Lab 2020 Pro software (Bruker). Principal component analysis (PCA) was used to compare different treatment groups to identify different ranks of variance within a data set *(Shao et al., 2012)*.

### Quantitative reverse transcription PCR (RT-qPCR)

Total RNA was isolated using an RNeasy Mini Kit (Qiagen). cDNA was synthesized using a RevertAid First Strand cDNA Synthesis Kit (Thermo Fisher Scientific). The generated cDNA was used as a template for RT-qPCR, which was performed with a ViiA™ 7 Real-Time PCR System (Applied Biosystems) using a PowerUp™ SYBR™ Green Master Mix (Thermo Fisher Scientific). PCR was carried out with an initial incubation at 50°C for 2 min, 95°C for 2 min, and then 40 cycles of 95°C for 15 s and 60°C for 1 min. The specificity of the reaction was verified by melt-curve analysis. The 2^−ΔΔCt^ method was employed to calculate relative gene expression levels. GAPDH was amplified as an internal control. The forward and reverse primers for GAPDH were 5’-GGTGAAGGTCGGTGTGAACGGA-3’ and 5’-TGTTAGTGGGGTCTCGCTCCTG-3’; for IL-1β 5’-GGAGAACCAAGCAACGACAAAATA-3’ and 5’-TGGGGAACTCTGCAGACTCAAAC-3’; for IL-2 5’-CCTGAGCAGGATGGAGAATTACAT-3’ and 5’-TCCAGAACATGCCGCAGAG-3’; for IL-6 5’-CCACTTCACAAGTCGGAGGCTTA-3’ and 5’-CCAGTTTGGTAGCATCCATCATTTC-3’; for IL-10 5’-GCCAGAGCCACATGCTCCTA-3’ and 5’-GATAAGGCTTGGCAACCCAAGTAA -3’; for CD86 5’-TCCTGTAGACGTGTTCCAGA-3’ and 5’-TGCTTAGACGTGCAGGTCAA-3’; for CD80 5’-GGTATTGCTGCCTTGCCGTT-3’ and 5’-TCCTCTGACACGTGAGCATC-3’; for CD40 5’-GTTTAAAGTCCCGGATGCGA-3’ and 5’-CTCAAGGCTATGCTGTCTGT-3’; for IL-9 5’-GTGACATACATCCTTGCCTC-3’ and 5’-GTGGTACAATCATCAGTTGGG-3’; for IL-10 5’-GCCAGAGCCACATGCTCCTA-3’ and 5’-GATAAGGCTTGGCAACCCAAGTAA -3’; and for TNF-α 5’-AAGCCTGTAGCCCACGTCGTA-3’ and 5’-AGGTACAACCCATCGGCTGG-3’.

### Statistical analysis

A Mann-Whitney *U* test or Student’s *t*-test was used to calculate statistical differences between different treatment conditions. Comparisons involving multiple groups were performed using 2-way ANOVA with Bonferroni’s multiple comparison test. The Kruskal-Wallis test, a non-parametric method, was used for MALDI-TOF data analysis. All the statistical analyses were calculated using Prism software 7. In all the statistical significance comparisons we have provided the exact *P*-values except for those *P-*values < 0.0001.

## Data availability

This study includes no data deposited in external repositories.

## Acknowledgements

We thank N.S. Li, Guangzhou Medical University, for technical assistance; C. Deng, C.M. Wong, Hong Kong Polytechnic University, for helpful advice on lipid metabolism; and K.L. Rock, University of Massachusetts, for providing the RF33.70 cell line. We thank the staff at the Centralized Animal Facilities (CAF) and University Research Facility in Life Sciences (ULS) at the Hong Kong Polytechnic University for valuable technical support. This work was supported by the Hong Kong Polytechnic University Internal Research Fund (P0001169), the NSFC/RGC Joint Research Scheme of the Research Grant Council of Hong Kong (N_PolyU533/19) and the National Natural Science Foundation of China (81961160738), PolyU Lean Launchpad Programme, and the Greater Bay Area International Institute for Innovations (GBA I3) Postdoc Programme.

## Author contributions

J-BS, JH and ZX conceived the study. ASC, J-BS, TJ, BW, JH, and ZX designed research. ASC, Y-WY, J-BS, and OZ performed research; All the authors contributed to data analysis. ASC, J-BS, JH, and ZX wrote the paper. All authors approved the final version of the manuscript.

## Supplementary Figures

**Supplementary Figure 1.**
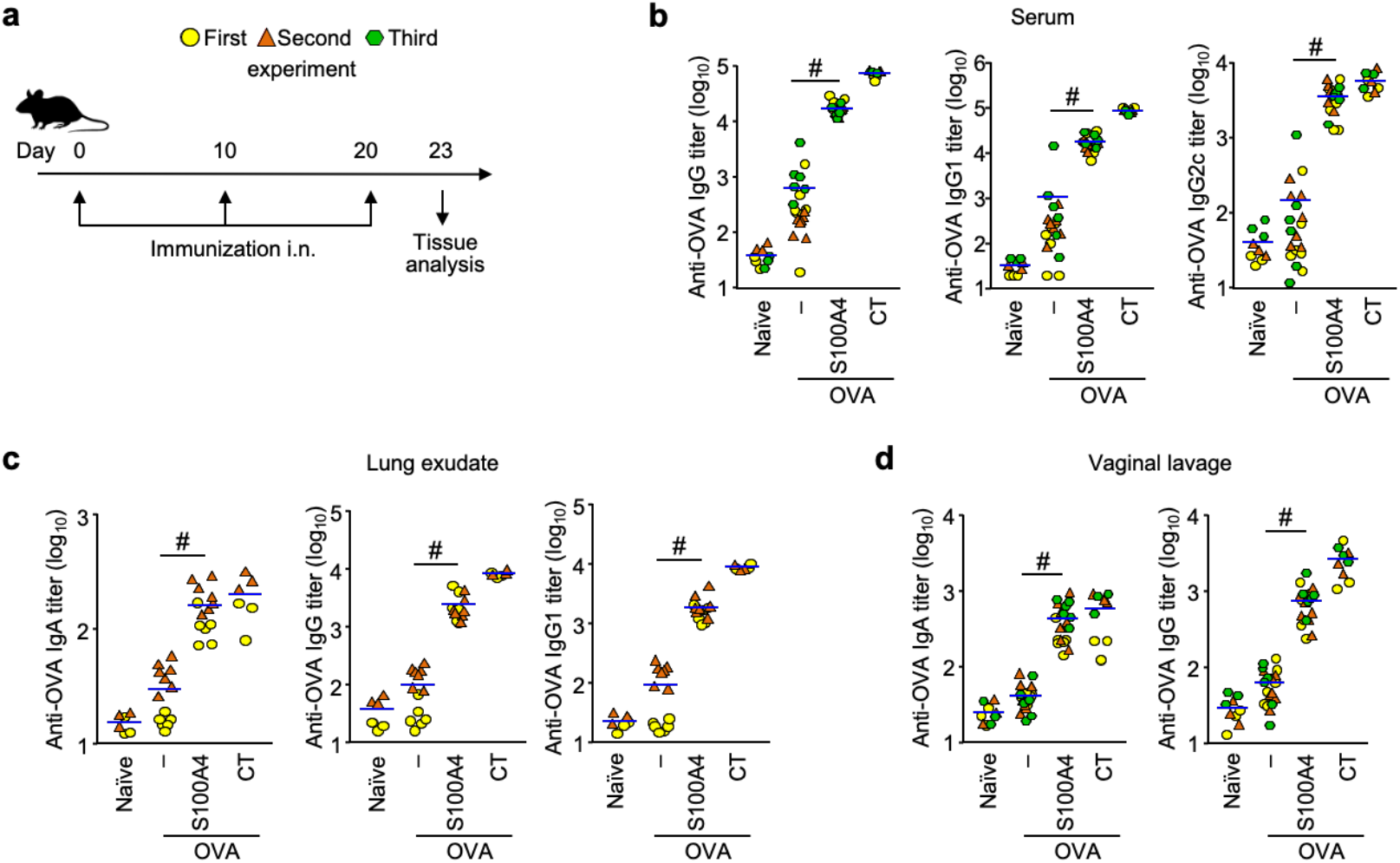
Intranasal immunization adjuvanted with S100A4 provokes greater humoral immune responses. Mice were immunized three times intranasally (i.n.) with ovalbumin (OVA; 10 μg) alone, or admixed to S100A4 (20 μg) or cholera toxin (CT; 1 μg) at a 10-day interval. Unmanipulated naïve mice were included for baseline control. Tissues were collected three days after the last immunization **(a)**. Various classes of OVA-specific antibody levels in serum **(b)**, lung exudates **(c)**, and vaginal lavage **(d)** were analyzed by ELISA. Each dot represents data from an individual mouse and blue lines indicate the average values. Symbols of the same colour represent data from one experiment. Data from three **(b, d)** or two **(c)** experiments were pooled. *#P* < 0.0001 by Mann-Whitney *U* test.

**Supplementary Figure 2.**
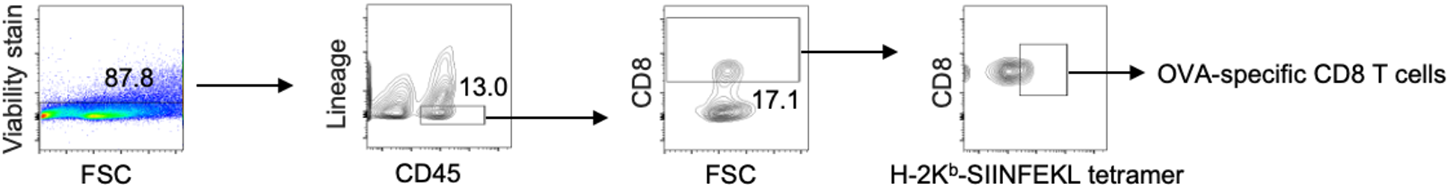
Gating strategies for lung CD8 T cell analysis are indicated. Mice were treated as explained in Figure 1 a. Lung tissues were harvested 196 days after the last immunization for flow cytometric analysis. A group of antibodies (Ter-119, GR-1, CD 11 b, CD11c and CD19) were used to exclude cells of unwanted lineages. Arrows indicate gating strategies in flow cytometric analysis. Numbers adjacent to outlined areas indicate percent cells in each gate. Shown are an example of a random lung sample. The gating strategy also applies to the investigation of the lung, lymph nodes and spleen at 10 days after the last dose of immunization.

**Supplementary Figure 3.**
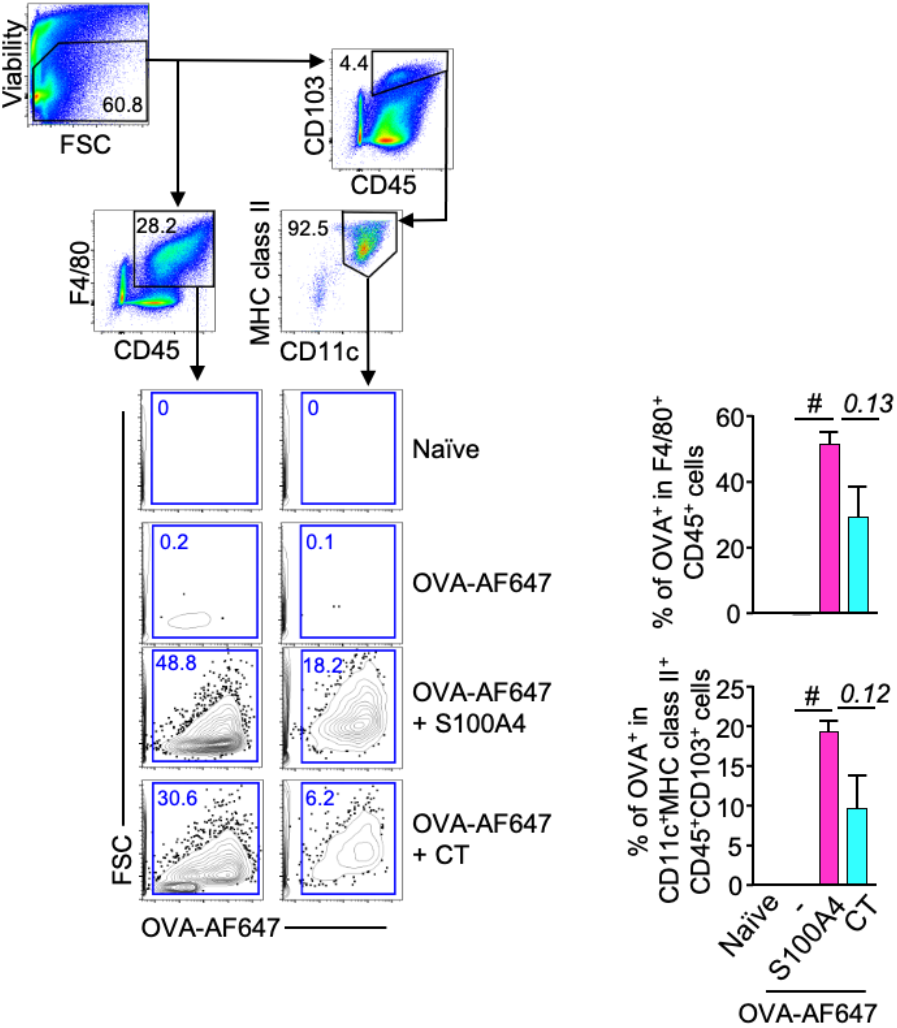
S100A4 promotes antigen transport to the lung. Mice received a single intranasal delivery of Alexa Fluor 647-conjugated OVA (OVA-AF647; 10 μg) in the absence or presence of S100A4 (20 μg) or cholera toxin (CT; 1 μg). Unmanipulated naïve mice were included for background control. Mice were euthanized at 12 hours after OVA administration and lungs were harvested for flow cytometric analysis. Representative contour plots indicating the biodistribution of OVA-AF647 in one representative mouse out of four in each treatment group are shown. Numbers in or adjacent to outlined areas indicate percent cells in each gate. Data are expressed as mean + s.e.m. of four biological replicates. *#P* < 0.0001 or the exact P-values (italic numbers) are indicated by unpaired *t* test.

**Supplementary Figure 4.**
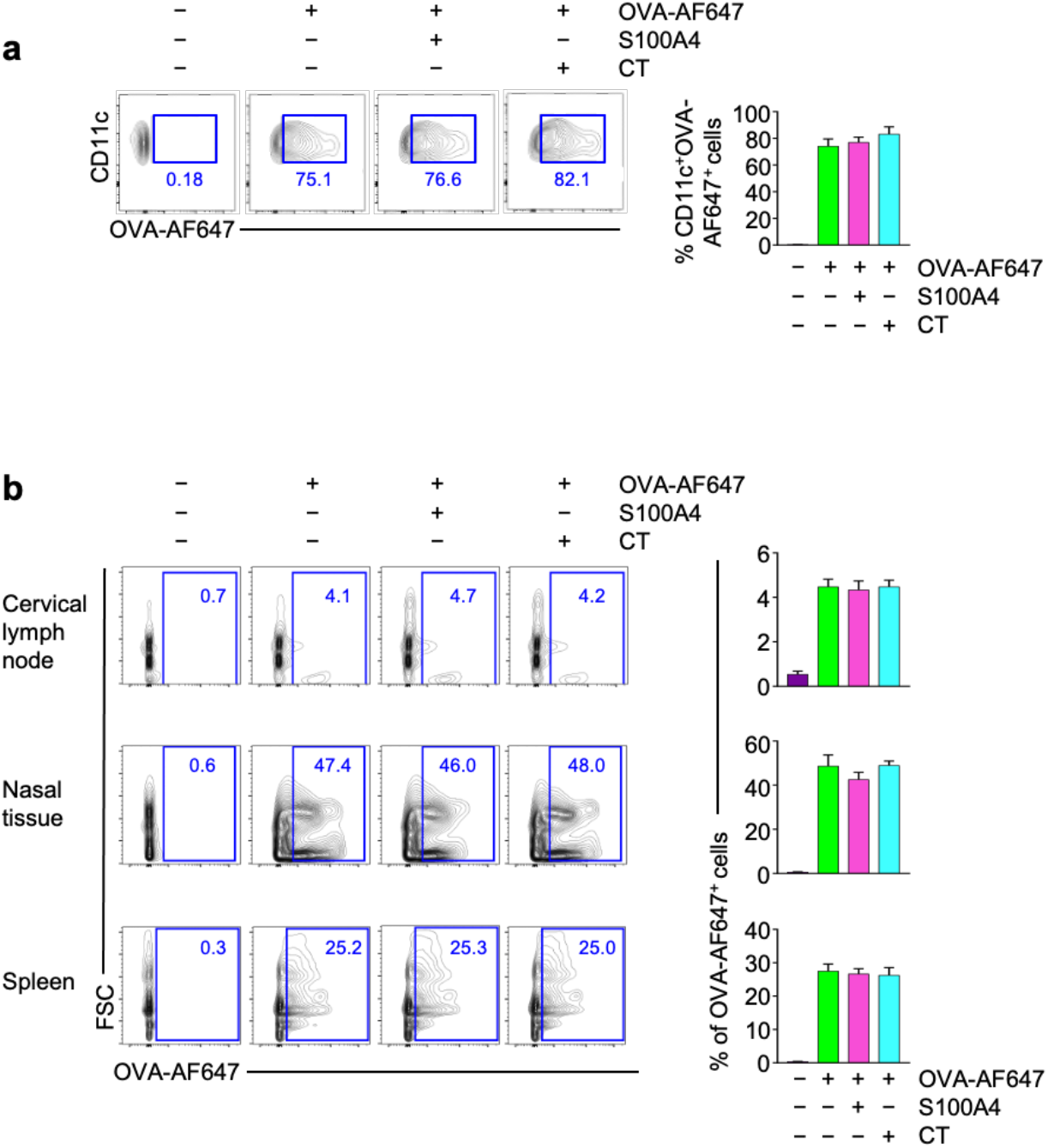
S100A4 does not promote antigen uptake by antigen presenting cells. Bone marrow-derived dendritic cells (BMDCs) **(a)** and mouse *ex vivo* cells from various tissues as indicated **{b)** were incubated overnight with or without Alexa Fluor 647-conjugated ovalbumin (OVA-AF647; 10 μg/ml), in the presence or absence of S100A4 (1 μg/ml) or cholera toxin {CT; 0.1 μg/ml), followed by flow cytometric analysis of the uptake of OVA-AF647. Representative flow cytometry contour plots {left panels) and quantification of the data (mean + s.e.m.) pooled from three mice (right panels) are shown. Numbers in or adjacent to outlined areas indicate percent cells in each gate. Shown is one of three similar experiments.

**Supplementary Figure 5.**
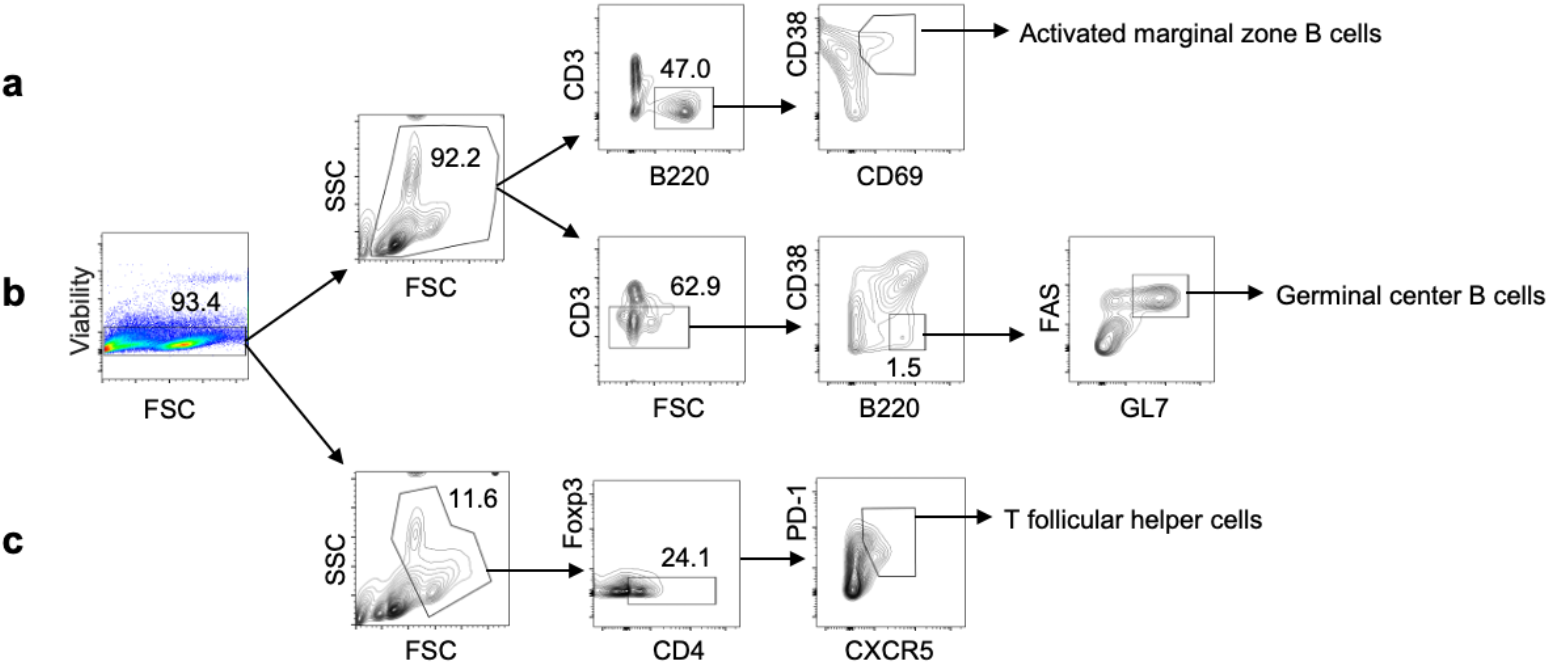
Gating strategies for splenic and lymph node cell analysis are shown. Mice were treated as in Figure EV1A. Spleens and cervical lymph nodes were harvested three days after the last immunization for flow cytometric analysis. Activated marginal zone B cells were identified as CD69^+^CD38^+^B220^+^CD3^−^cells **(a)**. Germinal centre B cells were identified as GL7^+^FAS^+^B220^+^CD3^−^CD38^−^cells **(b)**. T follicular helper cells were identified as PD-1^+^CXCR5^+^Foxp3^−^CD4^+^ cells **(c)**. Arrows indicate gating strategies in flow cytometric analysis. Numbers in or adjacent to outlined areas indicate percent cells in each gate. Shown are examples of a random spleen sample. The gating strategies and the contour patterns for lymph node analysis are similar.

**Supplementary Figure 6.**
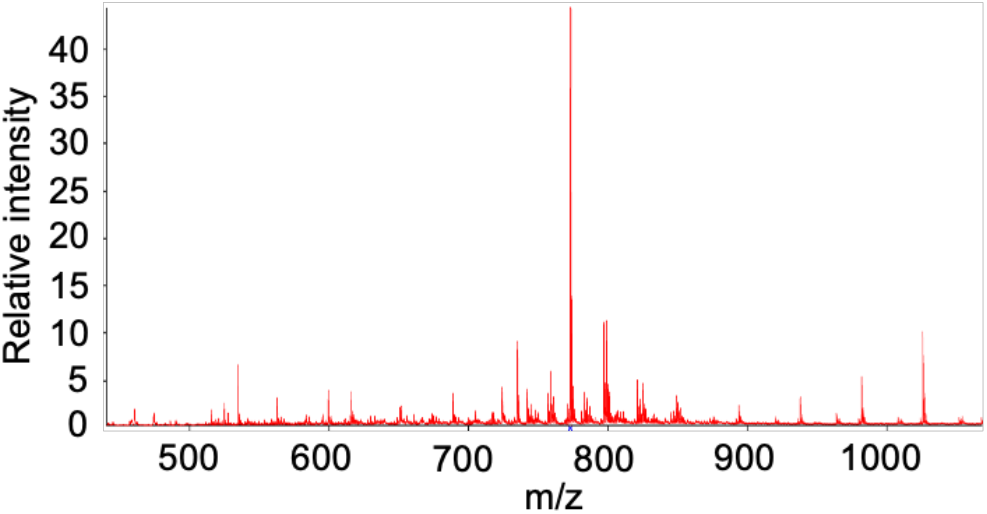
The mass spectra of all the lipids detected by MALDITOF are shown. Mice received three intranasal immunizations as explained in Supplementary Figure 1A. Spleens were collected three days after the last immunization and sectioned for MALDI-TOF analysis. A total of 348 lipids, including fatty acyls, glycerolipids, glycerophospholipids, polyketides, prenol lipids, and sterol lipids, were identified in the mouse spleens. About 83% of the total lipids were glycerolipids and glycerophospholipids. Shown is one example of a sample.

**Supplementary Figure 7.**
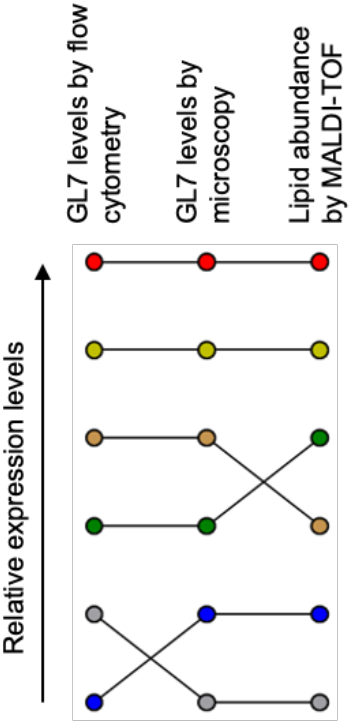
Spleen germinal center activities are consistent using different measurement techniques. Raw data displayed in Figure 4 and Figure 5 showing the expression levels of GL7 measured using flow cytometry or microscopy, as well as lipid abundance measured using MALDI-TOF, were processed and the expression levels of each (coloured dot) of the readouts were ranked at the single mouse level. Each line represents an individual mouse.

**Supplementary Figure 8.**
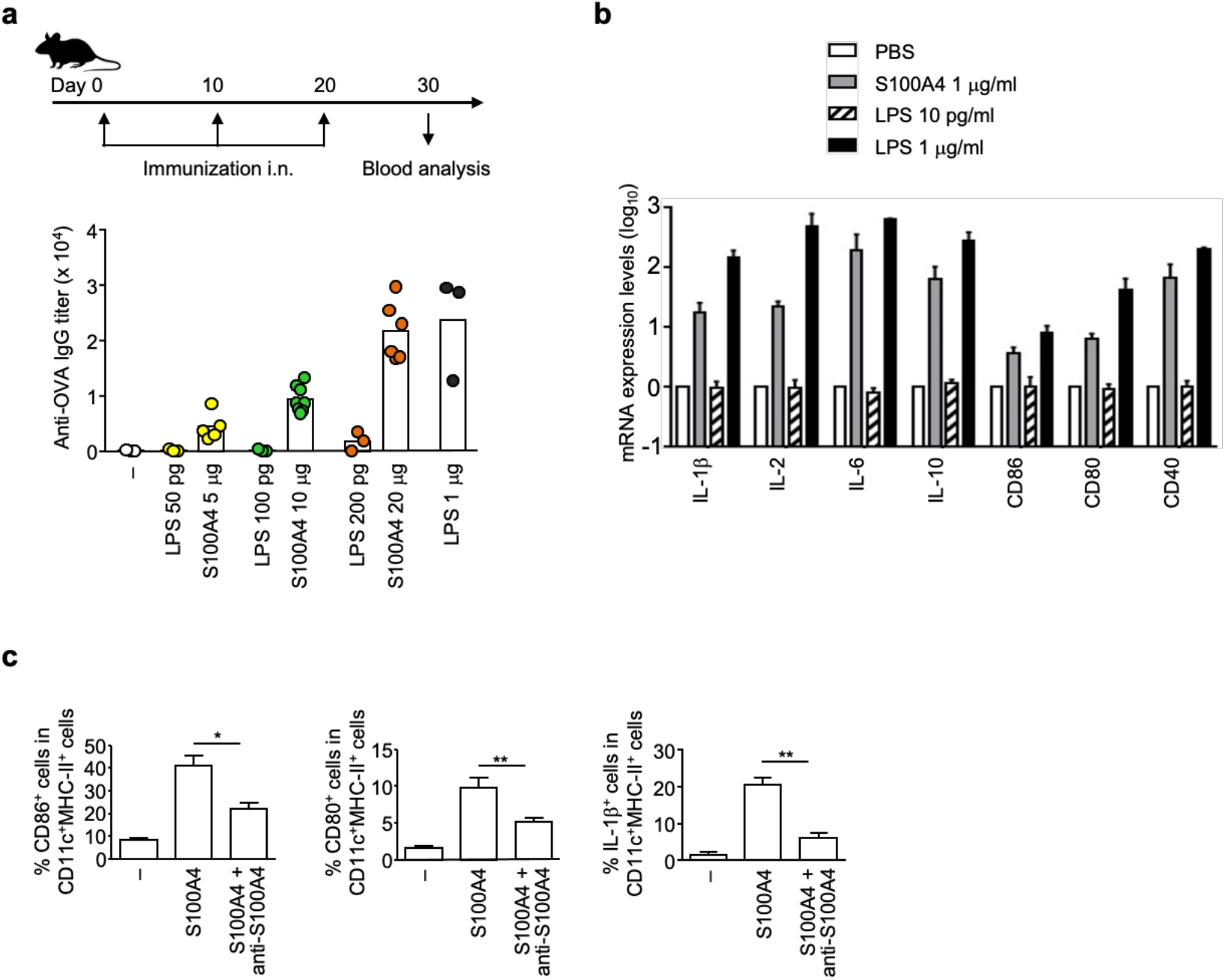
The adjuvant activity of S100A4 is not dependent on the residual amount of contaminating LPS. **a** Mice were immunized intranasally (**i**.n.) three times at a 10-day interval with ovalbumin (OVA; 10 μg) alone, OVA together with various amounts of S100A4 or with the corresponding residual LPS amounts present in each S100A4 dose. Serum was collected 10 days after the last immunization. OVA-specific total lgG levels were analysed by ELISA. Columns close to each other represent the pair that received an identical amount of LPS (exogenously added versus residual contamination in S100A4).**b** Bone marrow-derived dendritic cells (BMDCs) were incubated with S100A4 or the corresponding residual LPS amount present as indicated for three hours and levels of mRNA expression of a panel of cytokines that could be augmented by LPS were assessed using quantitative reverse transcription PCR. Gene expression was normalized using GAPDH as the calibrator gene. **c** BMDCs were incubated for one day in the absence or presence of S1 00A4, or S100A4 pre-mixed with an anti-S100A4 antibody. The frequencies of activated BMDC with enhanced expression of CD86, COBO or IL-1 *B* were measured using flow cytometry. Each dot represents data from an individual mouse and dots of the same colour indicate an identical amount of LPS the mouse received; columns indicate the average values **(a)**, or data are expressed as mean + s.e.m. of three biological replicates **(b)** or three separate experiments **(c)**. *^+^p* < 0.05; *^++^p* < 0.01 by Mann-Whitney *U* test.

**Supplementary Figure 9.**
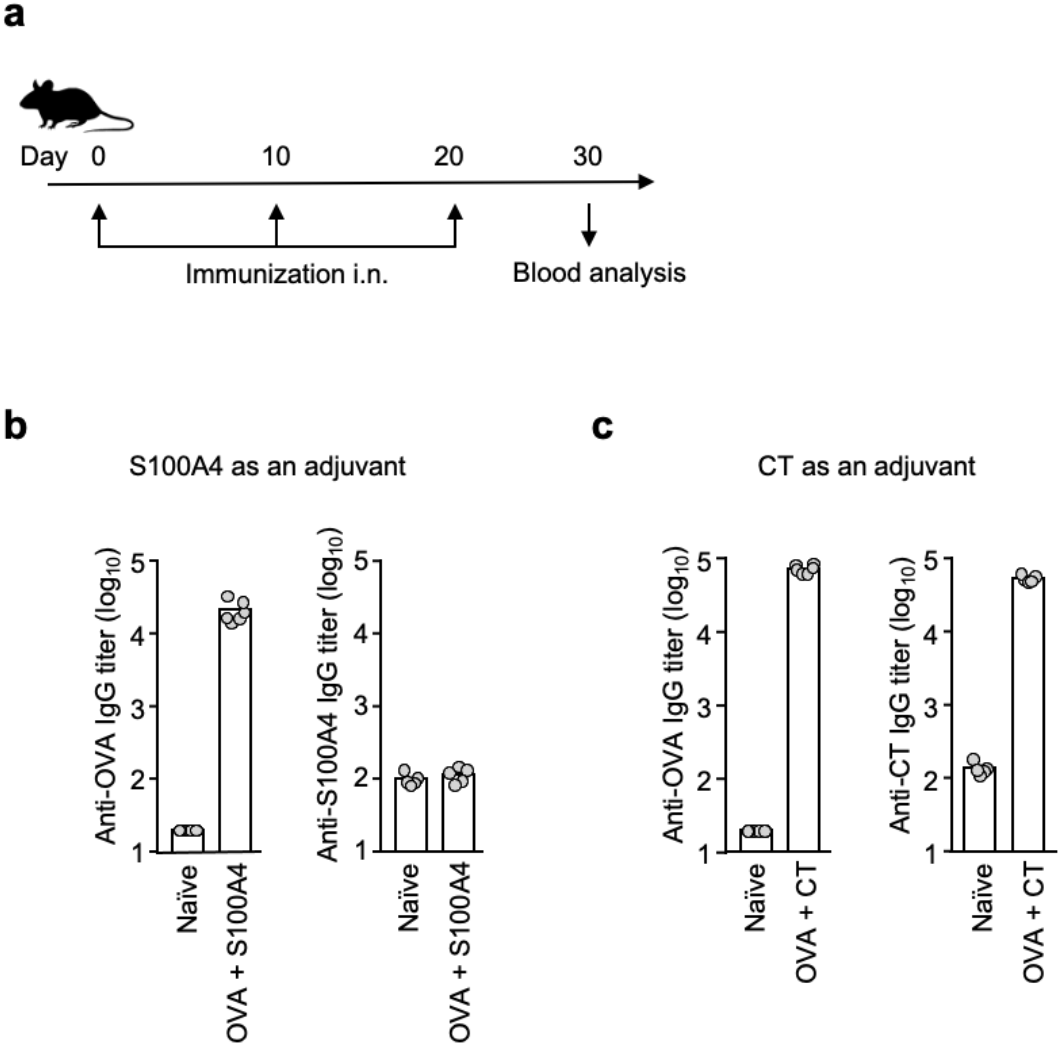
S100A4 does not induce anti-S100A4 antibodies after intranasal immunization. Mice were immunized three times intranasally (i.n.) with ovalbumin (OVA; 10 μg) alone or admixed to S100A4 (20 μg) or cholera toxin (CT; 1 μg) at a 10-day interval. Unmanipulated naïve mice were included for baseline control. Blood was collected 10 days after the last immunization **(a)**. Anti-OVA lgG antibody levels were determined for all the mice **(b, c)**. In addition, anti-S 1 00A4 and anti-CT lgG antibody levels were determined for those mice which received S 1 00A4 **(b)** or CT **(c)**, respectively. Antibodies were analyzed by ELISA. Each dot represents data from an individual mouse and columns indicate the average values.

